# Self-incompatibility limits sexual reproduction rather than environmental conditions in an invasive aquatic plant

**DOI:** 10.1101/2020.07.05.184267

**Authors:** Luis O. Portillo Lemus, Michel Bozec, Marilyne Harang, Julie Coudreuse, Jacques Haury, Solenn Stoeckel, Dominique Barloy

## Abstract

1. Fruitfulness and fertility are important components of sexual reproductive success in plants, and often depends on environmental conditions and reproductive systems. For invasive plants, fruitfulness and fertility control their ecological success and adaptation in invaded ecosystems. We studied which factors bring about fruitfulness and fertility in invasive populations of the aquatic plant *Ludwigia grandiflora* subsp. *hexapetala*.
2. We analysed fruitfulness and fertility of 37 populations growing under variable climatic conditions in Western Europe, and sub-sampled fruitful and fruitless populations grown in common controlled conditions. We carried out self- and cross-pollinations and measured their floral biometrics.
3. Environmental conditions, and temperature in particular, did not affect fruitfulness and fertility in-situ or in common controlled environments. Hand-pollinations resulted in fruit production by individuals sampled from fruitless populations when pollen came from fruitful populations, and by individuals sampled from fruitful populations whatever the origin of pollen. Floral biometrics evidenced the existence of two floral morphs that overlapped with fruitfulness, and individual incompatibility.
4. Our results rebutted the hypothesis that environmental conditions control fruitfulness and fertility in these invasive populations. We instead found that fruit and seed production were controlled by a reproductive system involving a self-incompatible approach herkogamous morph and a self-compatible reverse herkogamous morph. We assessed the floral morphs distribution worldwide of fruitless and fruitful native and invasive populations that matched our results at larger scale. Our results may constitute the first evidence of a possible heteromorphic self-incompatible system in *Ludwigia* populations and in Onagraceae phylogeny. It calls for further investigations on reproductive systems in this plant family. Finally, the observation that the self-incompatible morph seemed to be the world most invasive morph in this species tackles our understanding of biological and ecological conditions for invasiveness.
5. *Synthesis*. Our study showed that fruitfulness and fertility in the aquatic invasive plant, *Ludwigia grandiflora* subsp. *hexapetala* depend on a self-incompatibility system coinciding with two floral morphs, rather than environmental conditions and limitations. These new explanations on the sexual success of *Ludwigia* invasive populations will help defining new predictions about its worldwide spreads and ecological success, and will help reappraising future management plans.

## Introduction

Reproductive success is a central biological feature in understanding the ecology and evolution of population and species. It encompasses the ability of an individual or a population to produce descendants per breeding event, lifetime or generation (Rafferty, CaraDonna, Burkle, Iler, & Bronstein, 2013; Straka & Starzomski, 2015). In Angiosperms, the first two decisive steps of sexual reproductive success require that both the individual and population produce (i) fruit from their flowers by successful pollination and (ii) viable seeds from their gametes, which are able to germinate to give the next generation (Obeso, 2002). Fruitfulness (also referred to flower-fruit ratio, flower-to-fruit or fruit-set) is the ability of a plant to ensure fruit production from the pool of flowers to which a plant allocates its energy (Sutherland, 1986). Fertility is the ability of a plant to ensure the physiological maximum potential seed production from a gamete pool into their fruits (Aarssen & Taylor, 1992; Bradshaw & McMahon, 2008). Fruitfulness and fertility are essential for feed production and seed food as resources for ecosystems and human activities, and, as cornerstone components of reproductive success, they are essential for managing endangered, invasive, cultivated or unwanted populations (Barrett, Colautti, & Eckert, 2008).

Multiple environmental factors and intrinsic mechanisms may enhance or interfere with fruitfulness and fertility (Harder & Aizen, 2010; Sun et al., 2018; Sutherland, 1986). Environmental abiotic factors, such as sunshine, temperature and hygrometry, and biotic factors, including abundance of pollinators for entomophilous plants, flower grazers and seed diseases, affect plant fruitfulness and fertility (Giles, Pettersson, Carlsson-Graner, & Ingvarsson, 2006; Grass, Bohle, Tscharntke, & Westphal, 2018; McCall & Irwin, 2006). As a result, climate change could cause dramatic modifications in global angiosperm communities, with massive consequences for current ecosystem sustainability and agricultural production (Rafferty et al., 2013). Many intrinsic factors such as reproductive systems are key components to understanding population fruitfulness and fertility (Barrett, 1998; Dellaporta’ & Calderon-Urrea, 1993; Sutherland, 1986). The majority of flowering species present specific mechanisms for avoiding self-pollination, regrouped under the term of self-incompatibility (SI) systems, which result in the inability of individuals to produce zygotes after self-pollination (De Nettancourt, 2001). Self-incompatible species cannot self-pollinate, and therefore often present lower fruitfulness and fertility as a side effects, especially when populations lack compatible partners (Brys & Jacquemyn, 2020; Sutherland, 1986; Sutherland & Delph, 1984). Within SI species, heteromorphic SI systems include all plants with a physiological mechanism limiting self-pollination associated with two (distyly) or three (tristyly) different morphs of herkogamous flowers (Barrett, 2019). Herkogamous species present a spatial separation of their styles and anthers in hermaphrodites. Barrett (2019) defined two of herkogamy types: ‘approach herkogamy’, which corresponds to flowers with stigmas positioned above the anthers, and ‘reverse herkogamy’, for flowers with stigmas positioned below the anthers.

Distylous species genetically express two types of floral morphs, differing in their reciprocal heights of styles and stamen (reciprocal herkogamy). For example, in Primroses (*Primula veris & Primula vulgaris)*, Buckwheat (*Fagopyrum esculentum*) and Lungwort (*Pulmonaria officinalis*), individuals develop either long-styled or short-styled morphs, with short and long stamens respectively (Brys & Jacquemyn, 2009; Jacquemyn, Endels, Brys, Hermy, & Woodell, 2009; Li, Fang, Li, & Liu, 2017; Meeus, Brys, Honnay, & Jacquemyn, 2013). Tristylous species genetically express three types of floral morphs: long-styled, mid-styled and short-styled morphs, again differing in their stigma and anther/stamen heights. Distyly, the most-common heterostylous system was first documented by Darwin (1862), in the primrose (*Primula* spp.), which went on to become emblematic for this reproductive system (Gilmartin, 2015). Currently, distyly is described in 25 families, among which *Polygonaceae*, *Menyanthaceae*, and *Turneraceae* (Barrett, 1998). Tristyly, the rarest, has only been found in four families: *Oxalidaceae* (Turketti, Dreyer, & Esler, 2010), *Lythraceae* (Mal & Hermann, 2000), *Pontederiaceae* (Barrett, 1977; Glover & Barrett, 1983) and *Amaryllidaceae* (Barrett, 1998). Although these previous morphological features are valid for most heterostylous plants, several species show deviation from standard reciprocal herkogamy and morphologic compatibility patterns (Barrett, 1998). For example, *Narcissus assoanus* and *Jasminum malabaricum* present non reciprocal herkogamy, with only the style height dimorphism, while their stamens remain in the same position in both morphs (Cesaro & Thompson, 2004; Ganguly & Barua, 2020). In distylous, *Primula veris* and *P. vulgaris* show different rates of self-fertilization according to morphs (Brys & Jacquemyn, 2009; Jacquemyn et al., 2009), in *Luculia pinceana* and *Villarsia parnasiifolia,* only one of two morphs is self-compatible (Ornduff, 1988; Zhou, Barrett, Wang, & Li, 2015) and in distylous *Jasminum malabaricum* both morphs are self-compatibles (Ganguly & Barua, 2020).

The Onagraceae family includes about 657 species of herbs, shrubs, and trees in 17 genera (Les, 2017; Munz, 1942; Wagner, Hoch, & Raven, 2007; E. M. Zardini, Gu, & Raven, 1991). Figure S1 illustrates the wide floral diversity found in *Onagraceae* family. In this large family only the homomorphic gametophytic self-incompatibility (GSI) system has been described to date, based on studies on just two species: *Oenothera organensis* and *Oenothera rhombipetala* (Gibbs, 2014). Within Onagraceae, the *Ludwigia* genus includes 83 species, of which 75 species were classified as self-compatible, and 8 as self-incompatible (Table 3 (Raven, 1979)), which was then revised down a few years later to “*seven self-incompatible*” according to Zardini & Raven (1992).

Water primrose*, Ludwigia grandiflora* subsp. *hexapetala* (Hook. & Arn.) Nesom & Kartesz, (2000)), hereafter *L.g.hexapetala,* is a decaploid aquatic species which is one of the most aggressive aquatic invasive plants in the world (Thiébaut & Dutartre, 2009; Thouvenot, Haury, & Thiebaut, 2013). In the last decades, this species is reported as invading freshwater ecosystems in 15 countries worldwide, threatening local biodiversity, and water accessibility for human activities (EPPO, 2011; Hieda, Kaneko, Nakagawa, & Noma, 2020). Actions made by state- and privately-owned organisations aiming to limit the nuisance of invading *Ludwigia* spp. in only the lower part of the Loire basin in France cost around 340k€ in 2006 (Lambert, Genillon, Dutartre, & Haury, 2009). Historical data indicates that it originated in South America, and was introduced deliberately around 1826 in Europe (Dandelot, Verlaque, Dutartre, & Cazaubon, 2005). *L.g.hexapetala* populations reproduce using vegetative fragmentation and sexual seeds. Although its vegetative growth is particularly well documented, the role and importance of sexual reproduction within the population is still unknown, in particular how it contributes to the species invasiveness in newly-colonized areas (Dandelot, 2004; Ruaux, Greulich, Haury, & Berton, 2009; Thouvenot et al., 2013).

Invasive populations in France present contrasting fertility and fruitfulness depending on their geographical areas. Even though all populations massively bloom with the presence of a multitude of insects foraging flowers, the first appraisal of fruitfulness in *L.g.hexapetala* described fruitless populations in the Mediterranean zone, with flowers systematically degenerating after pollination, and fruitful populations yielding viable fruits and seeds on the European Atlantic coast (Dandelot, 2004). It led scientists and environmental managers to conclude that sexual reproductive success in invasive populations depends on climate factors (Dandelot et al., 2005). Invasive populations of *L.g.hexapetala* were described as having stigmas above the anthers on flowers with either 5- or 6 component parts in a distinct whorl of a plant structure (merosity) (Dandelot, 2004) and as putative allogame (Dandelot et al., 2005).

In this study, we thus aimed to identify whether climatic factors such as summer temperature and dryness affect the sexual success of the *L.g.hexapetala* in Western Europe, or if the mating system could explain the presence of fruitful and fruitless populations. Firstly, we tackled whether environmental factors really affect fruitfulness and fertility in those populations. To achieve this goal, we analysed i) fruitfulness and fertility in 37 *in situ* populations in varying environments, ii) fruitfulness and fertility of plants coming from fruitful and fruitless populations grown in a common garden with the same favourable environmental conditions. Secondly, we assessed this species regardless of temperature variation, by conducting hand-controlled pollinations on populations raised in a greenhouse, and measuring their resulting fruitfulness and fertility. Thirdly, since *L.g.hexapetala* showed variations in its floral morphology that can be interpreted as reverse herkogamy, we explored the sexual system in this species by studying the floral biometrics of different fruitfulness groups resulting from hand-controlled pollinations. We tested whether floral morphology *versus* temperature variation explain the fruitfulness variations measured in invasive populations in Western Europe. Finally, we discussed the non-implication of environmental factors on the sexual reproductive success of this invasive species, and the consequences of this first evidence of a heteromorphic self-incompatible system with two reciprocal-herkogames-morphs in the Onagraceae family.

## Material and Methods

### Studied species

*Ludwigia grandiflora* subsp. *hexapetala* [2n = 80] (Hook. & Arn.) Nesom & Kartesz (2000) populations are perennial in invaded areas (Dandelot, 2004; EPPO. 2011). Populations survive winter as submerged or buried plant parts. Rhizomes emerge and develop in dense mats from the spring to the following winter. Vegetative growth starts in April with the production of submerged foliage stems, then from June to early October foliage stems emerge, and flower buds are produced in each leaf axil (Thouvenot et al., 2013). Each stem produces a single flower every 3-4 days. Anthesis, the period during which a flower is fully open, with its pollen on open anthers and mucus visible on the stigma, occurs early in the morning (Dandelot, 2004). In the fruitful condition, three days after pollination, the floral receptacle, which contains the ovaries, remains green and continues to grow, whereas in the fruitless condition, the floral receptacle turns yellow and falls. Fruit (capsule) needs 6 weeks after pollination to become mature. *In situ*, bees, beetles and flies actively pollinate flowers. Without these pollinators, *L.g.hexapetala* remains totally fruitless, which leads to it being classified as strictly entomogamous (Dandelot et al., 2005). After pollination, fruit develops from August to November and the aerial parts of the plants degenerate in late autumn when seeds within the fruits are all mature.

### Sampling populations

We focused our study on populations invading Western Europe along a West-East transect in the Loire river watershed. The Loire receives the waters of one of the largest drainage basins in Western Europe and is the longest French river (Vogt et al., 2007). Its 117 500 km^2^ are known to cover a wide climatic gradient, from oceanic to continental, including variations in sunshine, temperature and precipitation. We monitored and observed the fruitfulness and fertility of 37 populations *in situ* over 765 km along Loire River (Table S1). We sampled individuals within 7 of these populations in order to study them in both common garden and greenhouse settings under controlled conditions (Table S1). To populate the common garden, we sampled 40 individuals per *in situ* population, extracted from five squares of 1 m^2^ at an interval of 10 m on a linear transect, in which we randomly chose 8 stems. All 40 individuals of a same population were installed together in a 450L mesocosm. A total of 280 sampled plants distributed in 7 mesocosms were installed in the common garden. Clonal cuttings of 10 randomly selected individuals per mesocosm were again installed together in 80L containers as replicates. This subsample of 70 plants distributed in 7 containers was installed in a greenhouse allowing us to manage the temperature (Schema of sampling protocol in Fig. S4). The common garden and greenhouse populations were installed in early June 2018 (location: Agrocampus Ouest, Rennes, France. 48°06′47.7″N 1°42′30.2″W). From May to October in 2018 and 2019, we watered both mesocosms and containers every 15 days with a commercial nutrient solution (NKP 6-6-6) during the growth and flowering periods, to ensure all plants were growing without nutritional deficiencies.

### Measurement of fruitfulness

In experimental populations, we measured the fruitfulness by comparing the number of flowers present and the quantity of fruit produced by the stem. In field populations, we measured the fruitfulness as the quantity of fruit produced per stem. Indeed, it has been previously documented that all populations in France massively bloom with the presence of a multitude of insects foraging the flowers (Dandelot, 2004; Ruaux et al., 2009; Thouvenot et al., 2013). Therefore, by only counting the number of fruits we assumed that all *in situ* populations have the same capacity for flower production. We named populations in which stems produced fruits as fruitful, and fruitless populations were those, which produced zero fruits. For experimental populations in the common garden, we counted flower and fruit productions on 40 randomly chosen stems in each mesocosm during the summer of 2018. For all 37 *in situ* and 7 common garden populations, we measured fruitfulness at the beginning of October as the number of fruits per stem produced on 40 randomly selected stems. For *in situ* populations, we delimited five 1 m^2^ squares on a 40 m transect separated by an interval of 10m, from which we randomly chose 8 stems and counted their fruit production. In greenhouse populations, we only measured fruitfulness on the hand-controlled pollinated flowers in each container. Three days after hand-controlled pollination, we counted the number of aborted flowers, and the number of fruits in formation. All fruits produced through controlled pollination were harvested at full maturity in order to assess their effective fertility.

### Measure of fertility

We measured fertility as the ability of fruit to produce seeds that successfully germinated and resulted in viable plants. For all fruitful populations among the 7 focused populations, we randomly sampled 5 ripe fruits per population both *in situ* and from the common garden, and 3 ripe fruits from the greenhouse. We counted the quantity of seeds produced by each of these ripe fruits (Fig. S6). We observed that success of seed germination in *L.g.hexapetala* required a seed dormancy interruption, but didn’t need vernalisation, i.e. floral induction by cold. We used a modified Hussner *et al.*, (2016) germination method: we put fruits in water in Petri dishes at 4 °C for a minimum of 3 weeks (cold-stratification period). We then deposited the seeds in basins in soil saturated with water at a temperature of 25 °C and a photoperiod of 16:8. Seeds began to germinate after 4 to 7 days. We measured fertility as the number of plants obtained one month after germination.

### Controlled pollination

In common garden populations, we isolated each mesocosm with an insect-proof net, only allowing intra-population pollination. To ensure pollen transport between and within flowers, we supplied mesocosms with ~ 350 pollinating flies (*Calliphora erythrocephala*) per mesocosm, every 3 weeks during the flowering period.

In greenhouse populations, we carried out two types of hand-controlled pollination: intra-individual pollination (self-pollination) and inter-individual pollinations, i.e. cross-pollination between individuals from the same population (named intra-population controlled cross) or from another population (named inter-population controlled cross). To perform hand-controlled pollination, when flower buds appeared, we locked them in cellophane bags to protect the flowers from incoming pollen. To ensure self-pollination, we shook the flowers in the bags after anthesis, and checked that pollen was deposited on the stigma. For inter-individual pollination, in order to simulate free random crosses, we selected five pollen-donor flowers and five other flowers which were to receive the selected pollen on their pistil. The pistil-donating flowers were emasculated before anthesis, and then pollinated with a mix of pollen from the 5 pollen-donating flowers. After pollination, the five pollinated flowers were sealed in cellophane bags to protect them from uncontrolled pollen incomings.

### Implication of climate and temperature in fruitfulness & fertility

To assess whether fruitfulness correlated with climatic conditions *in-situ*, we first compiled measures of sunshine, temperature and precipitation from the *Meteo France database* that occurred during the flowering time from June to August. We averaged climatic measurements that had been recorded every hour over the last 20 years (http://www.meteofrance.fr/climat-passe-et-futur/bilans-climatiques), and matched them with the fruitfulness measured in the 37 locations of *in situ* populations on Figure 1 and Table S1.

**Figure 1:**
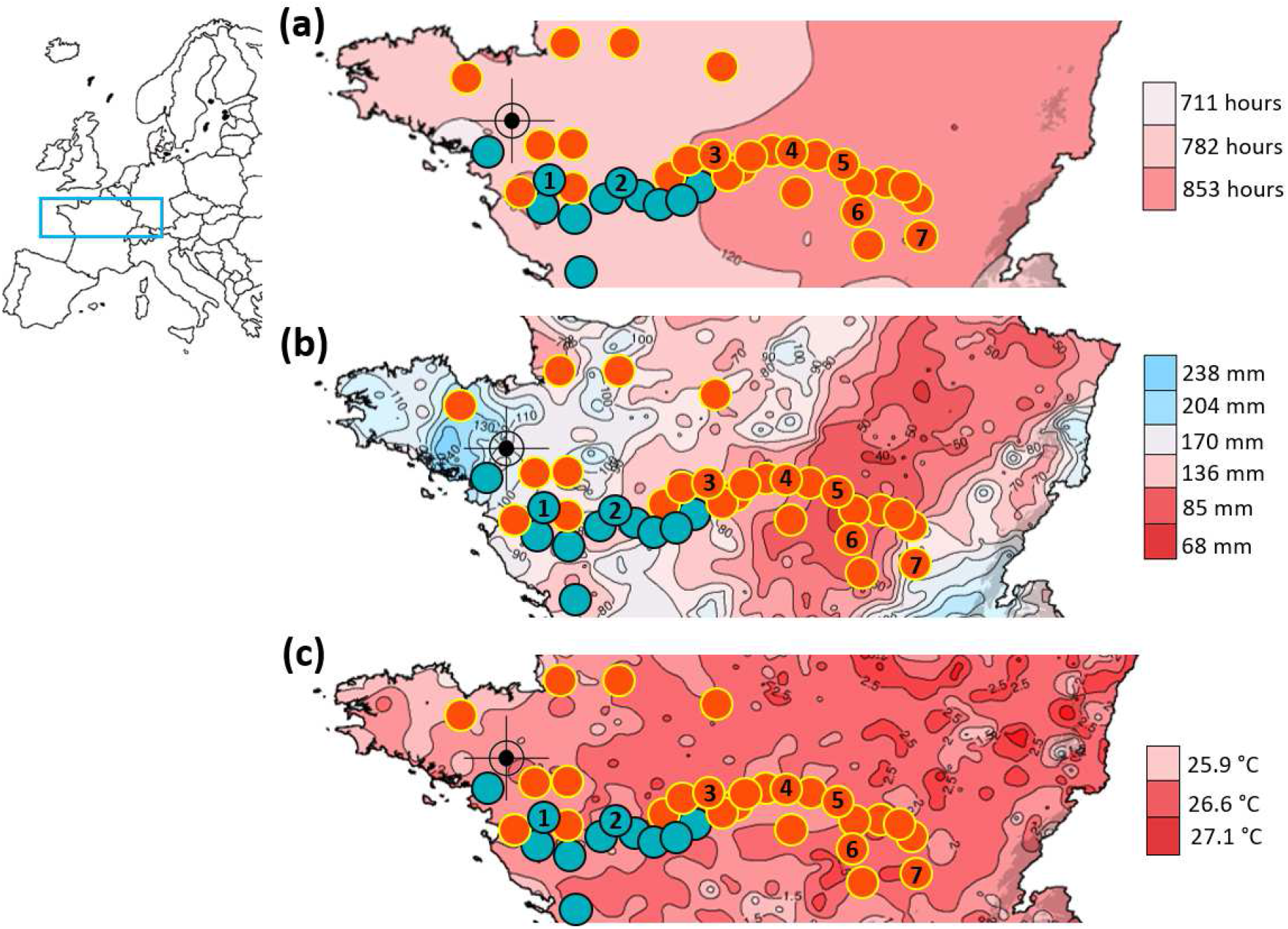
Location of 11 studied fruitful (blue circles) and 26 fruitless (orange circles) populations mapped with climatic conditions. Number in circles indicates the geographical positions of the 7 sampled populations: 1: Maze (Mazerolles); 2: Pont (Pont-de-Cé); 3: Orl (Orléans); 4: Poui (Pouilly-sur-Loire); 5: Gill (Gilly-sur-Loire); 6: Chat (Châtel-de-Neuvre) and 7: Cham (Chambéon) populations. The dark target symbol locates our common garden. See support information Table 1 for GPS Locations. (a) heat-map of cumulative sunshine hours in summer; (b) heat-map of cumulative millimetre of precipitation in summer; and (c) heat-map of the mean summer temperatures. All three maps were generated from the meteorological databases of “météo France”.

Secondly, to validate results obtained from *in situ* observations, we developed an experimental approach in the common garden, which ensured the same climatic and environmental conditions for all individuals sampled from different populations. If fruitfulness and fertility were controlled by environmental conditions, a common garden should homogenise plant fruitfulness and fertility, and their values would deviate from those observed *in situ*. We quantified the fruitfulness in the common garden populations in early October 2018 and their fertility in January 2019 alongside fruits sampled from field populations.

Thirdly, we carried out hand-controlled pollinations in the greenhouse under controlled temperatures. We performed the same replicates of self, intra-population and inter-population hand-controlled pollinations every 15 days from May to August 2019 at seven increasing temperatures (18° C, 22 °C, 27 °C, 30°C, 35°C, 40°C and 45° C, each varying by a max. +/− 2°C over a full day) to measure its impacts on fruitfulness and fertility within and between the seven sampled populations. At each temperature, we carried out 5 self-pollinations and 5 intra-population cross-pollinations per population; and 5 inter-population cross-pollinations for each of the 42 inter-population combinations. The fruitfulness data obtained in both *in situ* and experimental population observed were Bernoulli-type 0/1, and analysis was conducted using R software (R Core Team, 2014) using binomial generalized linear models (GLMs), with formula GLM (formula = Fruitfulness ~ “mean-of-summer-temperature” + “hours-of-sunshine-in-summer” + “summer-cumulative-precipitation”, family = binomial (link = “logit”)), and all figures in this paper were produced using the package ggplot2 (Wickham, 2009).

### Hand-controlled pollination between populations in the greenhouse

We assessed the sexual compatibility of individuals regardless of environmental conditions by conducting a full scheme of hand-controlled pollinations between populations growing in the greenhouse, and then measuring their fruitfulness and fertility.

From mid-July to early August 2018, we quantified the fruitfulness of individuals of (i) 105 self-pollinations, (ii) 105 intra-population pollinations, (iii) 630 inter-population pollinations crosses between individuals from different populations. Indeed, crosses of the 7 focused populations resulted in 42 inter-pop combinations. For example, we performed 15 cross pollinations with Maze-♀ × Pont-♂ and 15 cross pollinations with Maze-♂ × Pont-♀ (Fig. 3). In January 2019, we quantified the fertility of all fruit obtained from fruitful crosses.

### Floral morphology & biometrics

*L.g.hexapetala* showed variations in its floral morphology that can be interpreted as reverse-herkogamy. To explore its reproductive system, we thus studied floral biometrics on different fruitfulness groups resulting from hand-controlled pollinations.

Firstly, in all mesocosms of the common garden, from mid-July to early August 2018, we recorded the merosity of all flowers that blossomed on a total of 480 flowers for each fruitful or fruitless population, and computed the frequency distribution of merosity per population.

Secondly, at the same time, we measured 10 floral morphological traits on a total of 60 flowers. As *L.g.hexapetala* was previously described as 5-merous, we focused our measures on only 5-merous flowers. We measured morphological traits from 30 flowers sampled on individuals from fruitless populations (6 flowers per population × 5 fruitless populations) and 30 flowers sampled on individuals from fruitful populations (15 flowers per population × 2 fruitful populations). The 10 floral morphological traits that we measured were: length and width of sepal and petal, length of stamen and anther for the first and the second whorls, length of pistil and width of the floral receptacle (Table S2). A total of 300 floral parts (5 floral parts per flower × 30 flowers × fruitful & fruitless populations) were measured with a digital caliper (0.01 mm accuracy), except for styles. We measured 150 styles from 75 flowers from fruitful populations and fruitless populations, respectively. We also quantified nectar production on five flowers per population using a 10µL micropipette (Table S2).

All the floral parts measurements analysis was conducted using R software (R Core Team, 2014), using a Principal component analysis (PCA) and the Ade4 package (Dray & Siberchicot, 2020) to analyze correlation between floral morphological traits. To select the most discriminant variables we utilised a multiple regression analysis ((lm (Floral_groups ~ Pistil_length + Stamen.length_wh.1 + Stamen.length_wh.2 + Wh.2.Stament.pistil_ratio + Wh.1.Stamen.pistil_ratio + Sepale.length + Sepale.width + Petale.length + Petale.width + Anthere.length_wh.1 + Anthere.length_wh.2 + Receptacle.width) and the correlation figure was produced using the package “PerformanceAnalytics” (Table S4). AIC (Akaike information criterion) was utilised as a selection criterion.

## Results

### Environmental implication in fructification success

All the 37 monitored populations in the Loire basin massively blossomed from June to August. However, most of the populations (~70%, 26/37) were fruitless (Table S2). We mainly found fruitful populations on the ocean side, and fruitless populations on the continental side (Fig. 1 and Table S1). Climatic data showed variations in sunshine, cumulative precipitation and mean summer temperatures among studied populations (Fig.1). Cumulative sunshine hours ranged from 711 to 853h in two main areas: the first, eastern area only included fruitless populations, enclosing Mazerolles and Pont-de-Cé, while the western area encompassed both fruitful and fruitless populations, enclosing populations in Châtel-de-Neuvre and Chambéon (Fig. 1a). On the Atlantic coast, we found fruitful and fruitless populations with the same cumulative sunshine. Differences in pluviometry ranged from 86 mm to 204 mm, but again fruitful and fruitless populations were found in places with similar pluviometry on the western part of the watershed (Fig. 1b). Mean summer temperatures had little variation among populations, and ranged from 25.9 to 27.1°C. Again, fruitful and fruitless populations presented very similar values for this climatic factor (Fig. 1c). As expected, spatial distribution of populations on the Loire river depended on climate parameters with a clear differentiation between western populations on the first axis of PCA (R^2^=83.9%), which included fruitful Pont and Maze populations, and eastern populations as Chat, Cham (Fig. 1, Fig. S2). Principal Component Analysis also showed that chosen populations for the common garden experiment represented the climate variations measured on Loire River. Binomial generalized linear models (GLMs) analysis showed that the three environmental variables had a significant effect on fruitfulness. Temperature and precipitation showed negative correlation with fruitfulness (p-val= <2e^−16^), and sunshine showed positive correlation with fruitfulness (p-val= <2e^−16^). According to this result alone, a change in environment would lead to a change in fruitfulness, and thus several populations coming from different environments with contrasting fruitfulness should homogenize their fruitfulness in a common garden.

### Fruitfullness in the common garden

In the common garden with an identical environment (same climate, substrate, growth conditions and supply of pollinators), individuals sampled from the seven studied populations produced the same quantity of flowers per stem (p-value= 0.2998), between 6 to 15 flowers per stem regardless of the population origin (Fig. 2a). Only flowers from the two fruitful populations (Maze, Pont) produced fruits, between 6 and 15 capsules per stem, with all their flowers finally giving fruits (fruits-flowers ratio=1, Fig. 2b). Fruit production per stem was similar in Maze and Pont populations, whether *in situ* or in common garden samples (Fig. 2b, p-value= 0.2377). Samples from both fruitful populations however produced more fruit per stem in the common garden than monitored *in situ* (Fig. 2b, p-value= 0.0002). This result could be explained by the fact that fruits fall when they reach maturity. *In situ* fruit sets were only monitored once in October, whereas in the common garden they were monitored twice a week. By measuring the fruit production per stem in field populations in October, we may have missed early fruits that had already fallen.

**Figure 2:**
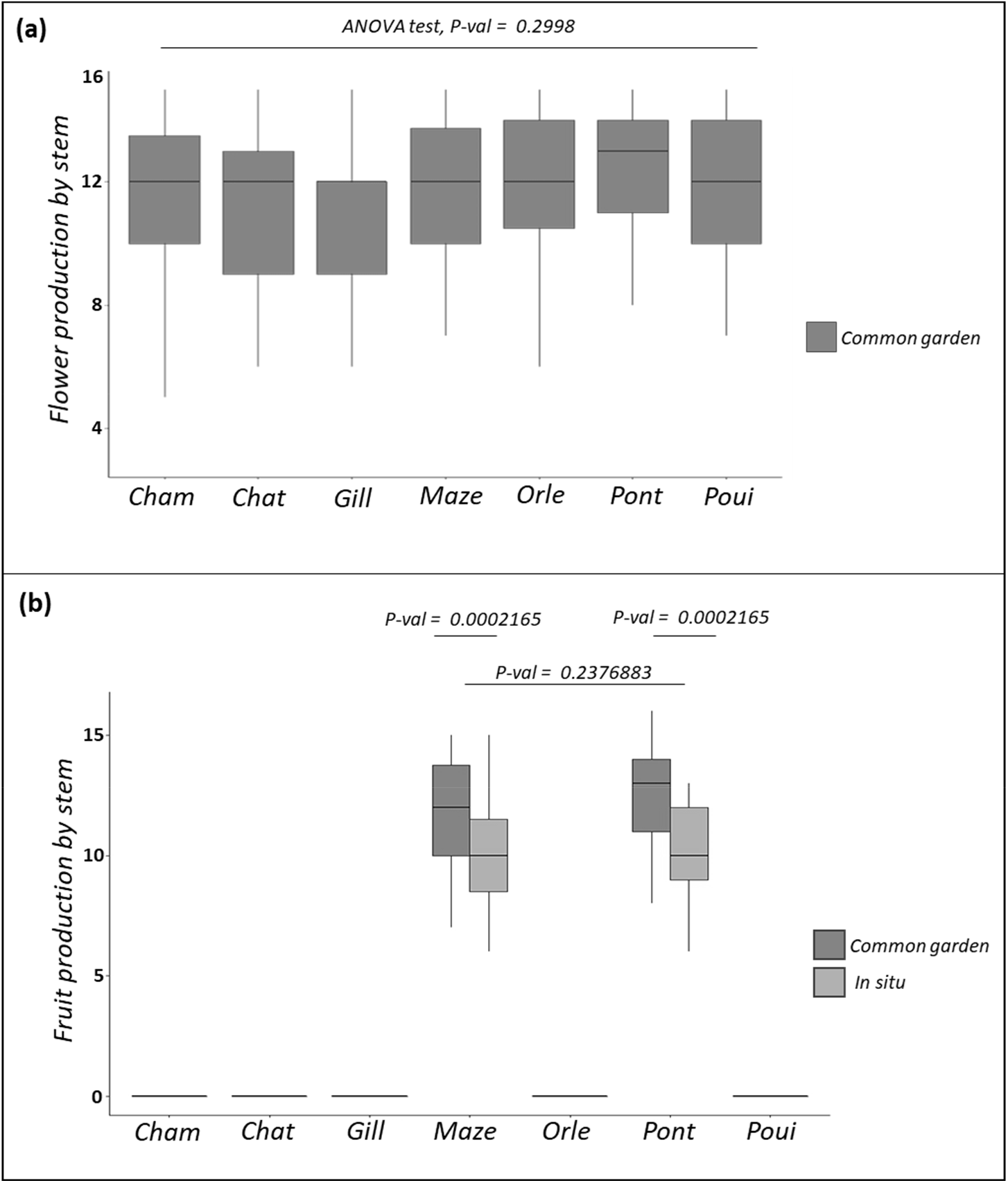
Distributions of flower (a) and fruit (b) productions per stem and population of L. grandiflora subsp hexapetala obtained in common garden (blue boxes) and in situ (green boxes) from July to early October 2018. (a) All experimental populations produced the same quantity of flowers in the common garden (p-val = 0.2998; Anova: lm (flower production ~ population)). (b) In both *in situ* and common garden, we observed two types of population: fruitless and fruitful. Maze and Pont were fruitful in both common garden and in situ conditions. The five populations Cham, Chat, Gill, Orle and Poui in both common garden and in situ conditions were fruitless. The two fruitful populations produced the same quantity of fruits *in situ* and in common garden (HDS test, p-val = 0.2376883), even though both of their fruit productions were higher in common garden conditions (HDS test, p-val = 0.0002165).

In the common garden, the 40 individuals sampled from one geographical population were free to pollinate one another. Despite sharing the same environmental conditions, individuals sampled from fruitful populations remained fruitful while individuals sampled from fruitless populations remained fruitless (Fig. 2b), indicating that environmental conditions didn’t affect fruitfulness.

### Fruitfulness from hand-controlled pollination

In the greenhouse, hand-controlled self-pollination and intra-population pollination gave the same results obtained in the common garden and field monitoring (Figs. 2, 3; Table S1). Flowers of individuals coming from the fruitful populations in Maze and Pont all produced fruit with a fruit-flower ratio of 1, while flowers of individuals coming from fruitless populations all became dehiscent and no fruit was produced (Fig. 3a, b). When flowers of individuals from fruitful populations were supplied with pollen from fruitless populations, the resulting fruit-flower ratio was still 1. Interestingly, supplying flowers of individuals from fruitless populations with pollen of fruitful populations resulted in a fruit-flower ratio of 1, and individuals from fruitless populations became fruitful (Fig. 3a, b).

**Figure 3:**
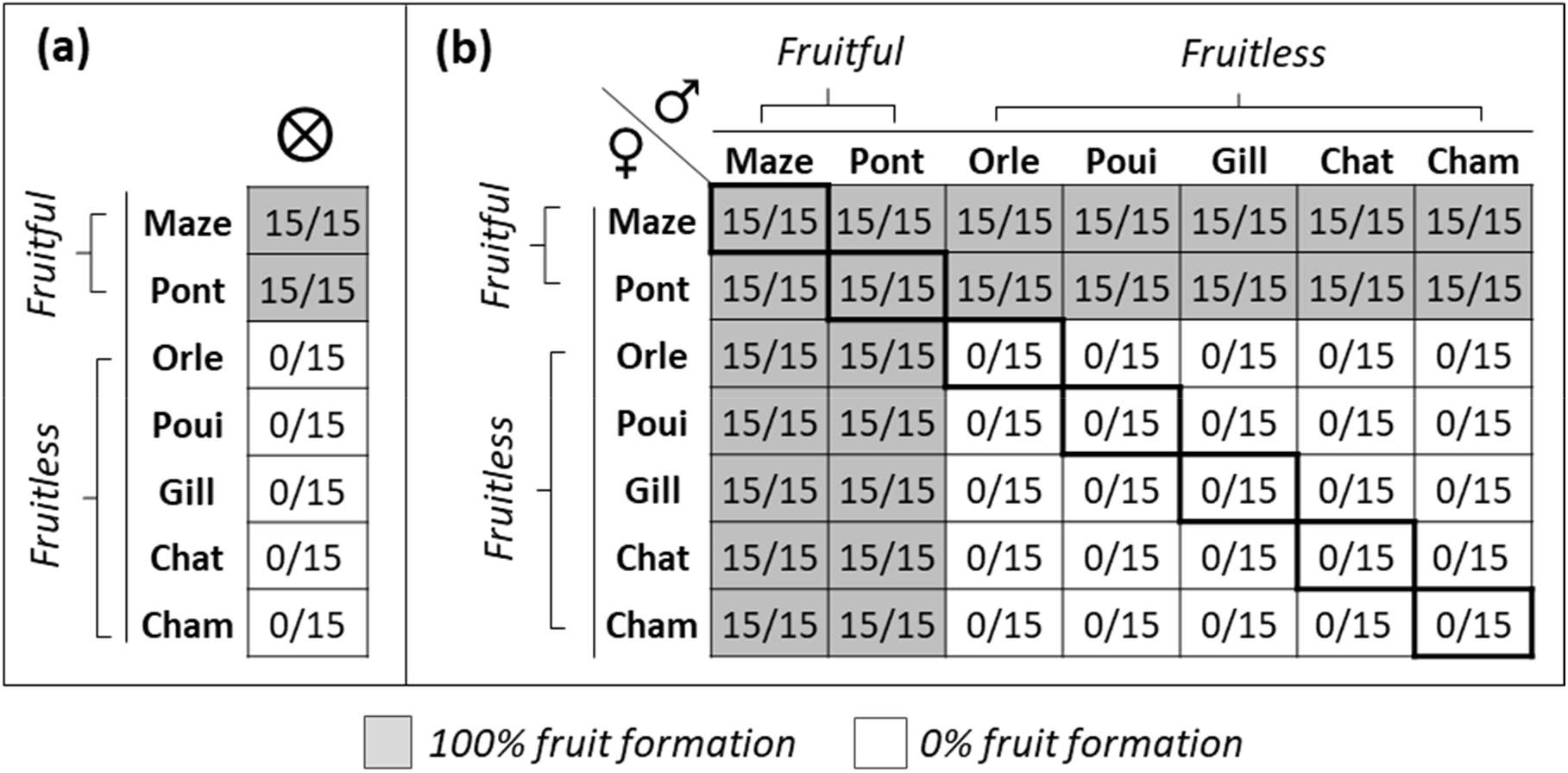
Fruitfulness in the 7 sampled populations of *Ludwigia grandiflora* subsp. *hexapetala* after hand-controlled-pollination crosses from mid-July to beginning of August 2018 in greenhouse. (a) Results of self-pollinations. (b) Results of intra-population (diagonal line) and inter-population (other boxes) hand-controlled cross-pollinations. Numbers separated by a slash indicate the ratio between fruits obtained from a fixed −15− number of flowers. A total of 840 pollinated flowers were hand-controlled pollinated: 105 self-pollinated flowers, 105 intra-population cross-pollinated flowers and 630 inter-population cross-pollinated flowers, corresponding to 15 flowers per population per pollination condition.

In 2019, hand-controlled pollination performed at 7 temperatures (18, 22, 27, 30, 35, 40 and 45°C) in the greenhouse showed congruent results (Fig. 4). Again, pollinating flowers of individuals sampled from fruitless populations with their own pollen or pollen from other fruitless populations resulted in no fruit at all temperatures (Fig. 4a, b). The same protocol applied to individuals sampled from fruitful populations resulted in a full fruit set, for all temperatures between 18°C and 45°C.

**Figure 4:**
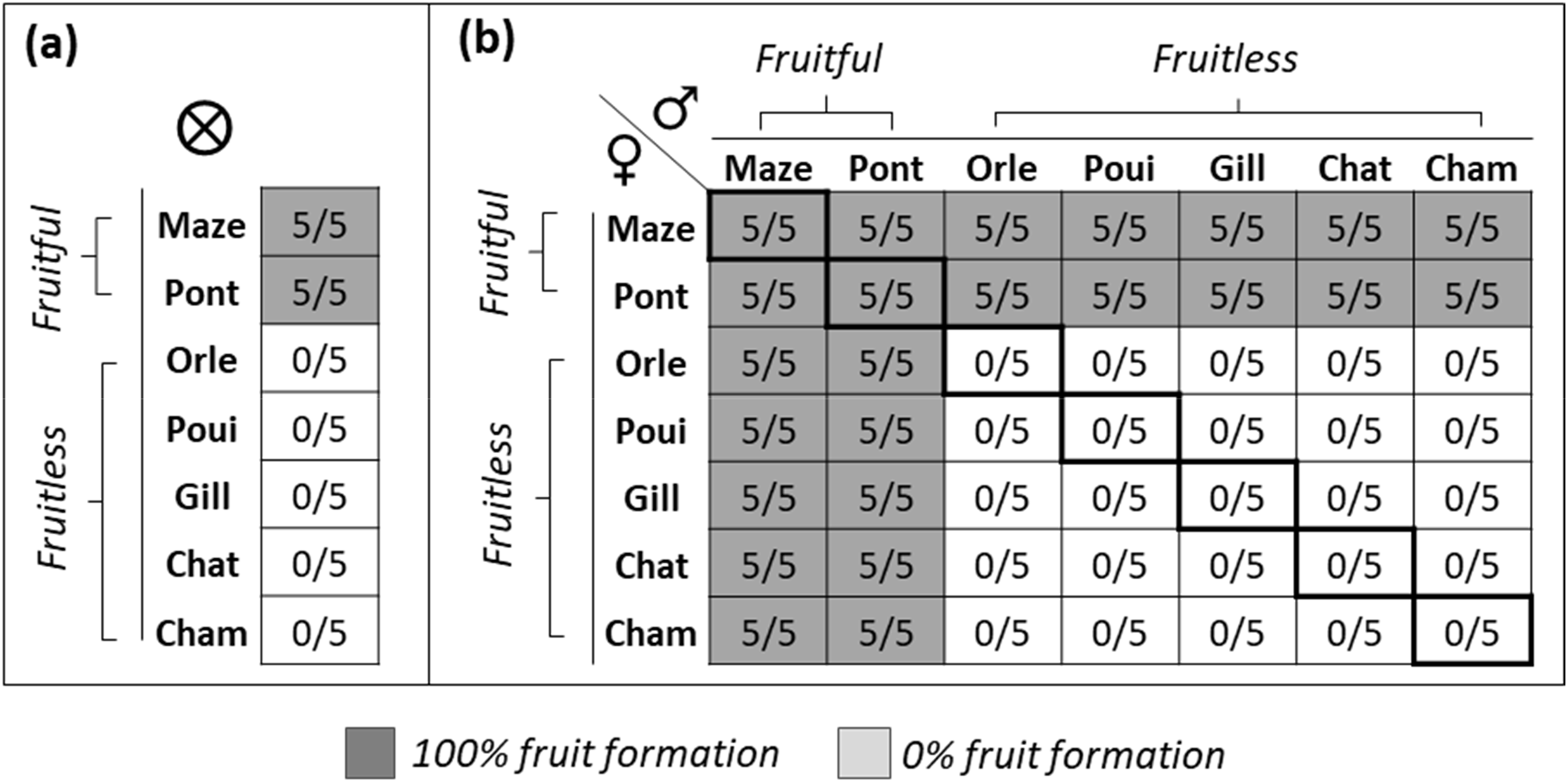
Fruitfulness in the 7 sampled populations of *Ludwigia grandiflora* subsp. *hexapetala* after hand-controlled-pollination crosses at 18 °C. Controlled-pollinations from May to August 2019 in greenhouse showed the same result on replicates performed under six different temperatures 22, 27, 30, 35, 40 and 45°C. We let the temperatures rise to its next higher value 15 days after performing the replicated hand-controlled pollination. (a) Results of self-pollinations. (b) Results of intra-population (diagonal line) and inter-population (other boxes) hand-controlled cross-pollinations. Numbers separated by a slash indicate the ratio between fruits obtained from a fixed −5− number of flowers for each temperature. Considering all temperate replicates, a total of 1960 pollinated flowers were observed = 245 self-pollinated flowers, 245 intra-population pollinated flowers and 1470 inter-populations pollinated flowers (5 flowers per population per condition and per temperature).

### Floral morphology in fruitful and fruitless populations

All populations produced 5, 6 and 7-merous flowers (Fig. S2). However, distribution of merosity type differed among populations. Both fruitful populations from Pont and Maze respectively produced 70 and 80% of 5-merous flowers, 25 and 15% of 6-merous flowers, and less than 5% of 7-merous flowers’ (Fig. S2a, b, c). Fruitless populations showed a differing distribution of merosity: 90 to 95% of 5-merous flowers, 4 to 9 % of 6-merous flowers and less than 1% of 7-merous flowers (Fig. S2d, e, f).

Measurements of floral parts on 30 flowers coming from fruitful and fruitless populations showed that (1) length and width of all parts of the perianth (sepals and petals), (2) Length of all parts of the androecium (first and second whorls carrying stamens and anthers) and (3) width of the floral receptacle, were all significantly bigger on flowers from fruitful populations than flowers from fruitless populations (Table S2). On the other hand, pistil length was significantly higher in flowers of fruitless populations than in flowers of fruitful populations. The principal component analysis (PCA) confirmed these differences showing a bimodal separation of the flowers belonging to fruitful (left) and fruitless (right) populations (Fig. 5c). The set of variables measured explained 80.1% of the variance. Moreover, the calculation of the length ratio between the stamen and pistil for the 1^st^ and 2^nd^ whorls for both fruitful and fruitless flowers revealed no difference in ratio for the 1^st^ whorl, but a significant difference for the 2^nd^ whorl. In flowers of fruitful populations, the length of the stamen in the 2^nd^ whorl was superior to the length of the pistil (Fig. 5e). Regression multiple analysis confirmed the PCA result (Fig. S5). The whorl-2-stamen-pistil ratio was the most discriminating characteristic (Fig. S5), but as all parameters were correlated, flowers of fruitful populations were significantly larger than flowers of fruitless populations for the majority of their floral parts, which may allow the fruitful floral morph to be distinguished from the fruitless by the human eye.

**Figure 5:**
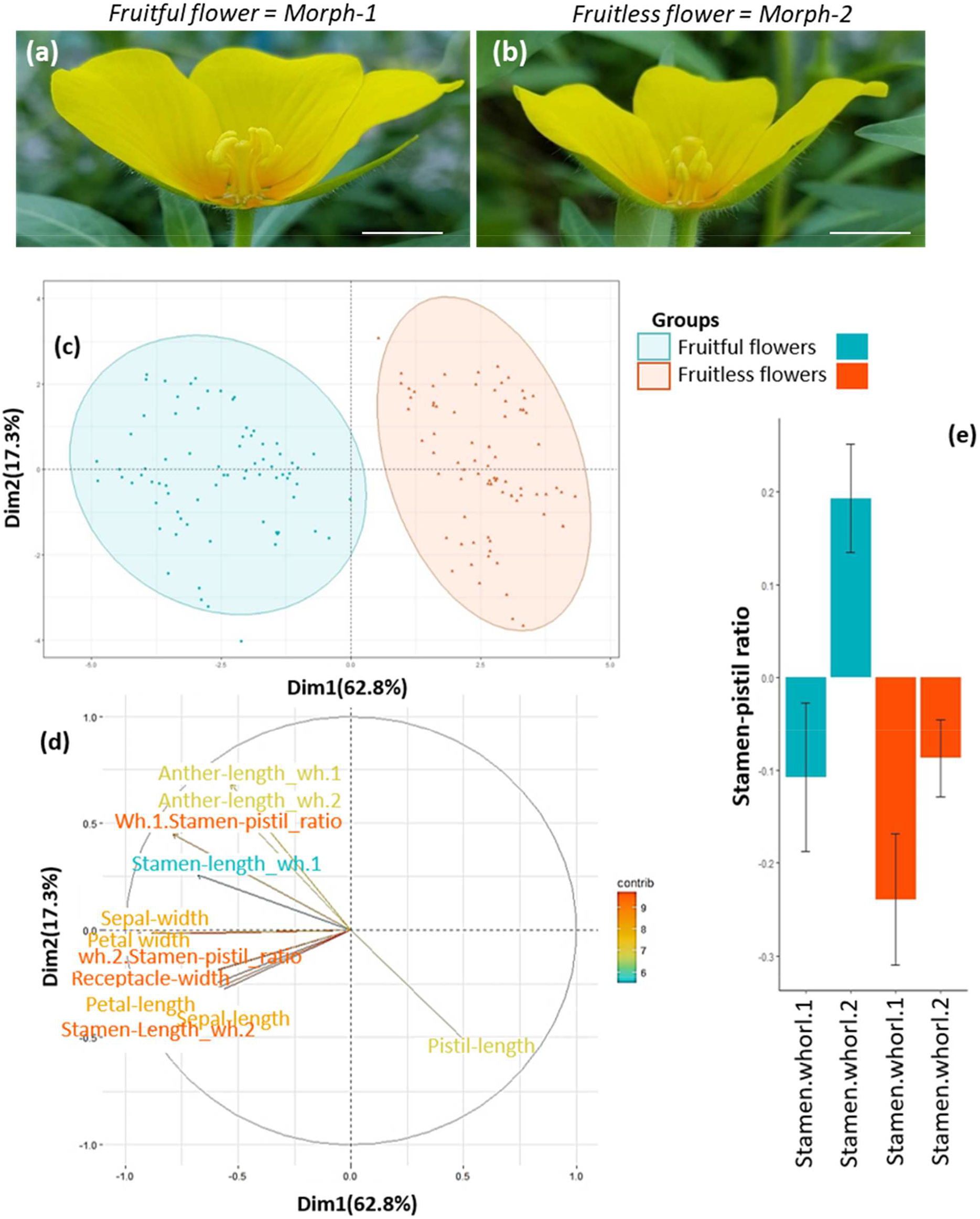
Floral biometrics of fruitful and fruitless populations in *Ludwigia grandiflora* subsp *hexapetala*. (a) Longitudinal sections of a typical flower from (a) fruitful populations and (b) from fruitless populations. Biometrics were measured from these sections. Bars = 1 cm. (c) Principal Component Analysis of floral biometrics: it showed two distinct clusters. These two floral groups fully coincided to fruitful (blue) and fruitless (red) groups (R2 =80.1%). (d): Variable factor map of floral biometrics. Colors indicated the variable contributions from low (blue) to high (red). For a given flower having a larger, perianth, androecium and floral receptacle showed a smaller pistil and vice versa. (Wh.1 or Wh.2= whorl.1 or Whorl.2)(e) Length stamen/pistil ratio of first and second whorls for fruitful (blue) and fruitless (red) flowers, bars represent standard deviations. In fruitful flowers, stamens from second whorl were always positioned above the pistil. In fruitless flowers, both stamen whorls were always positioned below the pistil.

This suggested the existence of two floral morphs with reciprocal herkogamy in *L.g.hexapetala*: The floral morph-1 (long stamined, reverse herkogamous flowers) was found in fruitful populations and the floral morph-2 (short stamined, approach herkogamous flowers) was only found in fruitless populations. *In situ* populations showed perfect congruence between their fruitfulness and the morphs of their flowers. In fruitful populations including those sampled in Maze and Pont, we only found individuals with morph-1 flowers. In fruitless populations, including those sampled in Orle, Poui, Gill, Chât and Cham, we only found individuals with morph-2 flowers.

### Linking fruitfulness of hand-controlled pollination with floral morphology

After self-pollination, all morph-1 flowers produced fruits while all morph-2 flowers were aborted (Fig. 6a). Pollinations between two different individuals of morph-1 flowers (morph-1 × morph-1; Fig. 6b) always produced fruits; conversely, pollinations between two different individuals of morph-2 were always unsuccessful (morph-2 × morph-2; Fig. 6b). Pollination between individuals with different morphs always produced fruits; independently of which morph was used as male and female plant (morph-1 × morph-2; morph-2 × morph-1; Fig. 6b).

**Figure 6:**
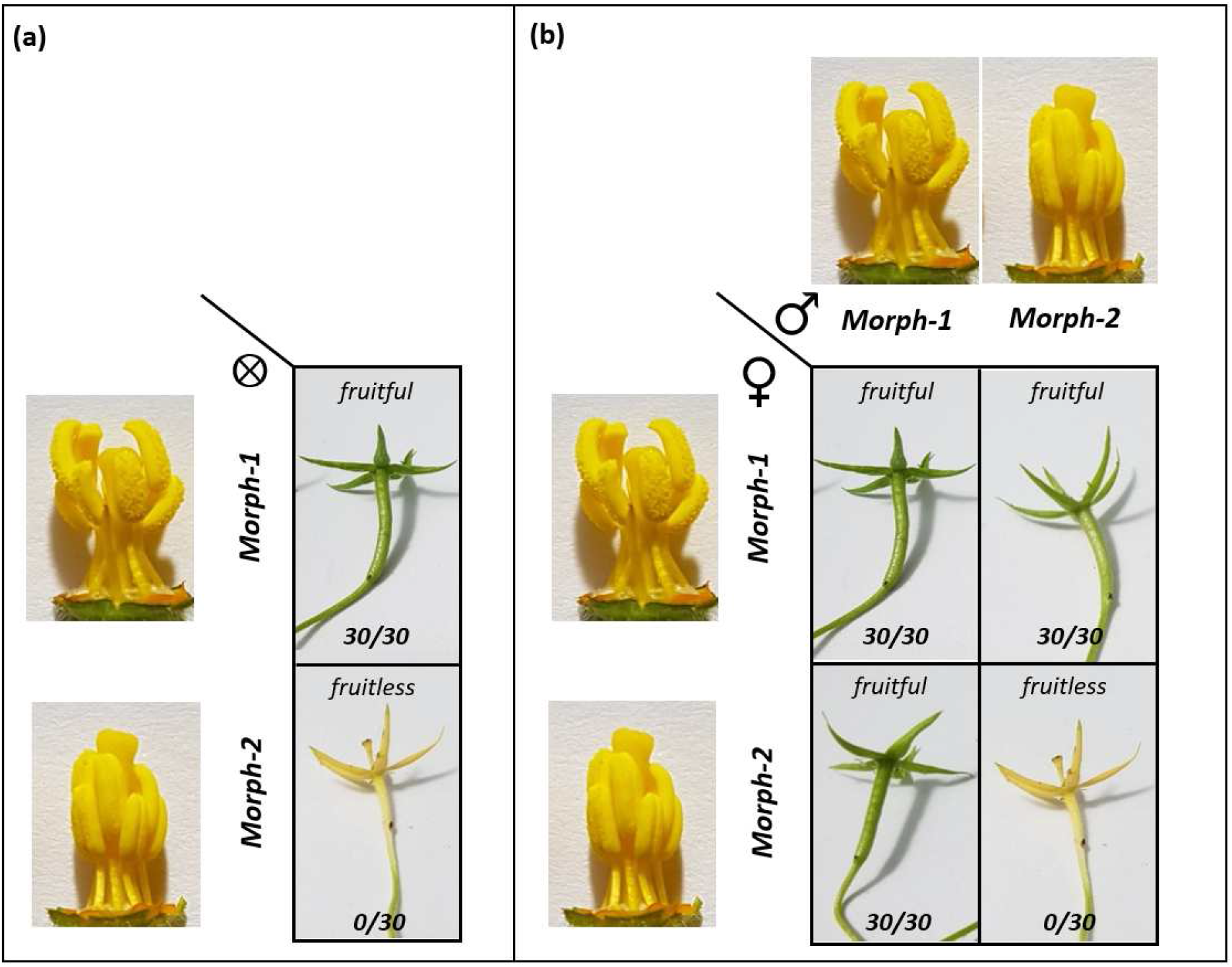
Fruitfulness in function of floral morphs and types of crosses in *L. grandiflora* subsp *hexapetala*. (a) Result of self-pollination of morph-1 or morph-2. (b) Result of intra-morph and reciprocal inter-morph crosses between morph-1 (fruitful populations) and morph-2 (fruitless populations). Green fruit were fruits in formation with developing seeds (fruitful); yellow flowers were dehiscent flowers with no fruit formation and no seeds (fruitless). Numbers separated by a slash indicate the ratio between fruits obtained from a fixed −30− number of flowers. In self-pollination and intra-morph cross-pollination, only morph-1 populations produced fruits. In inter-morph cross-pollination, all crosses with morph-1 used as male or female gave fruit.

We then assessed the viability of the seeds obtained from all these crosses. All the produced fruits contained between 59-72 fully-formed seeds, with the germination rate always superior to 90% (Fig. S6), regardless of the pollination condition (free entomophilous or hand-controlled pollination), population and floral morphology.

## Discussion

Our results argued that geographical distribution of a self-incompatible morph, rather than biotic or abiotic environmental conditions, explained the success of sexual reproduction of invasive populations of *L.g.hexapetala* in Western Europe. Our results also argued for the first evidence of a self-incompatibility system coinciding with two different floral morphs in this worldwide invasive species. if it were to be confirmed by additional studies, our results would constitute the first evidence in Ludwigia genus and in Onagraceae family for a heteromorphic self-incompatibility system.

### No environmental and/or climate factors are involved in the reproduction success of *L. grandiflora* subsp. *hexapetala*

Temperature was believed to be the main factor affecting *L.g.hexapetala* fruitfulness and fertility (Dandelot et al., 2005). Indeed, increases or decreases in temperature after flower induction in some other plant species may deteriorate the production of viable male gametes and cause male sterility (Liu, Mo, Zhang, De Storme, & Geelen, 2019; Santiago & Sharkey, 2019). In France, fruitful populations were initially observed in the Atlantic zone, while fruitless populations were found in the Mediterranean zone, hypothesizing a climate effect on the reproductive success of *L.g.hexapetala* (Dandelot et al., 2005). Our study, which focused on the Loire basin, confirmed the presence of fruitful populations in the Atlantic area, but also showed the presence of fruitless populations in this area. Moreover, we found no effect of temperature on fruitfulness between fruitless and fruitful populations. Common garden and greenhouse experimental populations produced viable fruits and seeds from 18 to 45°C. These results reject the hypothesis that the temperature may deteriorate and limit sexual reproduction in invasive *L.g.hexapetala*. The experimentations on fruitfulness and fertility performed in our common garden also showed that other environmental variations found between fruitful and fruitless populations in Western Europe cannot be used to explain their current fruitfulness and fertility. Indeed, all individuals maintained their fruitful or fruitless state when in common environmental conditions, and pollinated by individuals of the same population, while all populations produced the same quantity of fruits and viable seeds when supplied with compatible pollen.

### First evidence of a self-incompatibility system found in *L. grandiflora* subsp. *hexapetala*

Our hand-controlled cross-pollinations showed that *L.g.hexapetala* presented both self-compatible and self-incompatible populations in Western Europe. Haury, Damien, Maisonneuve, & Bottner, (2012) reported few cases of fruitless invasive populations becoming fruitful for the first time, when studying populations in Western France. After evidencing the self-incompatibility system coinciding with two floral morphs, we more recently found that 5 of our monitored (but not sampled for common garden experimentations) *in situ* populations observed to be fruitful resulted in fact in a mixture of individuals producing morph-1 flowers and individuals producing morph-2 flowers. In early October, in these populations, all individuals produced fruits regardless of their floral morph under free pollination. The coexistence of one compatible and one self-incompatible type of the same very invasive species raises many ecological and evolutionary questions, and highlights the necessity to carry out improved and further investigations into its sexual reproductive system. The first step will be to characterize the type of incompatibility system, i.e. sporophytic or gametophytic system through pollen germination and ovule fertility in both morphs, as has been done for *Guettarda speciosa* (Xu, Luo, Gao, & Zhang, 2018) and *Primula oreodoxa* (Yuan et al., 2018).

Raven (1979) studied the mating system in Onagraceae and classified the breeding systems of all 674 species: 283 (42%) are classified as outcrossing; 353 (52%) as self-pollinating and less than 6% (38) have a mixed breeding system. Among the 80 species of Ludwigia genera, 26, 54 and 0 species were classified as outcrossing, self-pollinating and mixed breeding systems, respectively. Our results revealed that *L.g.hexapetala* was not only strict allogamous, as hypothesized by Dandelot et al., (2005) but also reproduced using a mixed mating system relying on a self-incompatible system coinciding with two floral morphs. Les (2017) classified all North American *Ludwigia* species as mainly self-compatible, with only 8 putatively self-incompatible species in South America. Our identification of a self-incompatible floral morph and a self-compatible floral morph in *L.g.hexapetala* thus calls for a search for similar self-incompatible systems in other *Ludwigia spp.* and Onagraceae in general.

### Floral morphs are associated with sexual reproductive success in *Ludwigia grandiflora* subsp. *hexapetala*

We found that fruitful and fruitless populations of *L.g.hexapetala* showed different floral morphologies and merosity. In the *Ludwigia* genus, variations in merosity were already reported between species (Wagner et al., 2007). Here, for the first time in the *Ludwigia* genus, we showed that merosity variations occurred between and within populations of a single species, *L.g.hexapetala,* and its distribution may be linked to its floral morphs. It questions the ecological and evolutionary importance of such biological features in this genus.

Beyond merosity, the analysis of floral biometrics allowed us to highlight the existence of two flower morphologies for each fruitfulness population type. Barrett (2019) defined distyly as a genetic polymorphism where two mating types differ reciprocally in stigma and anther height. Our analysis of floral biometry showed two reciprocal herkogam morphs, with a ratio stamen-pistil greater than 1.25 and of less than 0.9, matching the most important Barrett’s (2019) criteria for functional herkogamy, and the pistils of the morph-1 flowers always 1-2 mm smaller than the pistils of the morph-2, also matching a secondary criteria of heteromorphic system (Barrett 2019). The floral characteristics we found in *L.g.hexapetala* corresponded to two other well-known distylous species: *Fagopyrum esculentum* (Li et al., 2017) and *Linum suffruticosum* (Ruiz-Martín et al., 2018). Both of these species show a non-tubular dystilous flower structure, we also found in *L.g.hexapetala* (Fig. S7). Altogether, our floral biometrics and cross-pollination experimentation may suggest an heteromorphic self-incompatible system in invasive populations of *L.g.hexapetala*. An old and abundant literature discussed floral morphology in Onanagracea, particularly in the *Ludwigia* species (Eyde, 1977; Raven, 1979). But, to our knowledge, floral dimorphism has been never mentioned in this family and genera before. We supposed that those morphological criteria might have been too subtle to be distinguished by eye and without dedicated measures. In addition to floral morphology, the main functional characteristic of heteromorphic systems is its assortative incompatibility, implying that all morphs are expected to be self and intra-morph incompatible (Barrett, 2019). However, several species have already been listed where one of the morphs does not follow this rule, or the morphs show different rates of self-compatibility (Brys & Jacquemyn, 2009; Ganguly & Barua, 2020; Ornduff, 1988; Zhou et al., 2015).

For *L.g.hexapetala,* we observed that morph-1 produced fruit after self- or intra-morph pollinations, which were never the case for morph-2. The only reciprocal inter-morph pollination which produced seeds was when morph-2 received pollen. These results suggested that (1) morph-1 was self and cross-compatible (intra and inter-morph), and (2) morph-2 was self-incompatible and intra-morph incompatible and only inter-morph compatible. Similar results have been observed in the distylous *Luculia pinceana,* with the L-morph self-compatible, the S-morph self-incompatible and both morphs intra-morph compatible (Zhou et al., 2015). We therefore call for further investigations in other invasive and native populations of *L.g.hexapetala* to strengthen or reject the hypothesis of an heteromorphic self-incompatible system in this species and assess the stability and reproducibility of the two very distinct floral morphs we evidenced.

### Fruitfulness and floral morphs association in worldwide populations of *L.g.hexapetala*

To confirm the existence of two morphs in other worldwide populations and confirm or reject their association with fruitfulness, we collected web data (from sourced photographs, herbaria, papers, wildlife services and surveys) on populations in native and invasive areas, to which we associated floral morphs (using our floral biometrics) with their reported fruit and seed productions (Table S3). Historical and recent data showed the presence of both self-incompatible and self-compatible types and floral morphs in the native area of *L.g.hexapetala* (Argentina, Southern Brazil, Uruguay) as well as in invaded areas (North America, Europe). Populations in which we only detected morph-2 flowers were always congruent with populations being reported as fruitless. Mapping the biogeography of fruitful and fruitless populations of *L.g.hexapetala* in native and invaded areas with self-incompatible and - compatible morphs would help future studies in terms of the understanding of genetic diversity, ecology, and evolution of this species and will allow to trace the timing and the routes of its spread worldwide, identifying vectors and characteristics of favourable environments.

### Self and inter-morph compatible system calls for increased management effort on fruitful populations

*L.g.hexapetala* is known as one of the most threatening invasive species worldwide. Its wide range of environmental tolerance on fruitfulness and fertility may partly explain its worldwide invasiveness, and managers should not uniquely consider environmental conditions or climate changes when trying to limit its expansion and proliferation. Modelling of *L.g.hexapetala* dispersal in terms of the climate predicts that its spread should increase up to 2 fold in Europe and North America by Gillard, Grewell, Deleu, & Thiébaut, (2017). Suitable new areas will mainly be located to the north of its current range.

However, we showed that fertility and fruitfulness were not affected by temperature in Western Europe, and sexual reproduction in these areas may exacerbate its expansion and proliferation, which should be considered in future plans for worldwide control. Indeed, it is only a matter of time before fruitless populations meet an incoming compatible morph and thus become fruitful.

Sexual reproduction in *L.g.hexapetala* may impact its invasiveness, as it decisively participates in its dispersal. Indeed, floating seeds present a greater dispersal distance than clonal fragments, over 1000km using water flow (Ruaux et al., 2009) and transport by vertebrates (García-Álvarez et al., 2015). The presence and persistence of *L.g.hexapetala* sexual seeds in seed banks (Grewell, Gillard, Futrell, & Castillo, 2019) highlight the importance of considering sexual reproduction in the resilience of this species when devising management plans. We roughly quantified that seed production in capsules was about 50000 seeds/m^2^. Current management plans in invaded areas mainly rely on clonal propagation (Dandelot, 2004). The demonstration that the temporal lack of compatible pollen suspended seed production is a definite game-changer in all strategies defined to control this species. In contradiction with Baker’s conjecture (Pannell, 2015), populations at the forefront of the invasion, in the populations we monitored in Loire basin and in the worldwide database we analysed (Europe, North America and Asia), were most often due to morph-2, the self-incompatible morph. On top of this, sexual reproduction with massive recombination can also generate new abilities and favour local adaptations through new genetic and epigenetic combinations, which can then be locally maintained by clonal reproduction. Interestingly, one of the first populations able to reproduce sexually in the European Atlantic area newly present unusual adaptation to the terrestrial environment through genetic and epigenetic factors (Billet, Genitoni, Bozec, Renault, & Barloy, 2018; Genitoni et al., 2020). To limit the risk of the appearance and dispersal of new genotypes, and indirectly avoid a secondary invasion, management recommendations should pay particular attention to fruitful populations, and regulate seed production, for example by preferentially planning elimination actions at the beginning of blooming to limit fruit and seed production.

In conclusion, we rebutted that environmental conditions, with a particular focus on a wide range of temperature, limited sexual reproduction in invasive populations of *L.g.hexapetala* as conjectured by previous literature and management plans (Dandelot, 2004). We also reported the first evidence of a self-incompatible system coinciding with two floral morphs in invasive populations of *L.g.hexapetala* in Western Europe which evokes a heteromorphic, dystilous, self-incompatible system. If confirmed by further studies, it would constitute the first evidence of a heteromorphic incompatible system in Onagraceae, with two floral morphs: one approach herkogamous self-incompatible and one reverse herkogamous self-compatible. An improved characterization of its heteromorphic incompatibility system in its physiological mechanism, and its genetics, should help us to understand its ecology and evolution, especially in invaded areas, and thereby be used to rationalize management plans. As a major invasive species with reproduction including self-pollination, allogamy and clonality, *L.g.hexapetala* presents interesting features in order to study the link between reproductive modes, success in species expansion, and adaptation to new environments, as tackled by Baker (1955). Finally, our study calls for a reappraisal of sexual reproduction in the *Ludwigia* genus and other Onagraceae genera, and provide some future elements to discuss in terms of the importance of reproductive modes in helping to shape angiosperm phylogeny.

## Acknowledgements

This research was supported by FEDER funds from Région Centre-Val de Loire and by Agence de l’eau Loire-Bretagne (grant Nature 2045, programme 9025 (AP 2015 9025)). FEDER also financed the doctoral grant of L. Portillo and the technical assistance salary of M. Harang. The authors thank Diane Corbin (FRAPNA Loire - Ecopôle du Forez), Guillaume Le Roux (Reserve Naturel Val d’Alier Châtel-de-Neuvre) for making plant material available and Romain Deschamps (Conservatoire d’espaces naturels de l’Allier) for the photos of Onagraceae species. We thank the Experimental Unit of Aquatic Ecology and Ecotoxicology (U3E) 1036, Institut national de recherche pour l’agriculture, l’alimentation et l’environnement (INRAE, which is part of the research infrastructure Analysis and Experimentations on Ecosystems-France, for help with the maintenance of plants. We thank Pascale Breger (Agrocampus Ouest INRAE, UMR SAS) for help to use of meteo France database. We thank S. Barrett for all the exchanges and comments on early results.

## Author Contribution

LP and DB designed this project. MB, MH, LP and DB performed all experiments. JC, JH and SS participated capsule husking. LP, SS and DB analysed data and wrote the manuscript. All authors read the manuscript.

## Supporting Information

**Figure S1:**
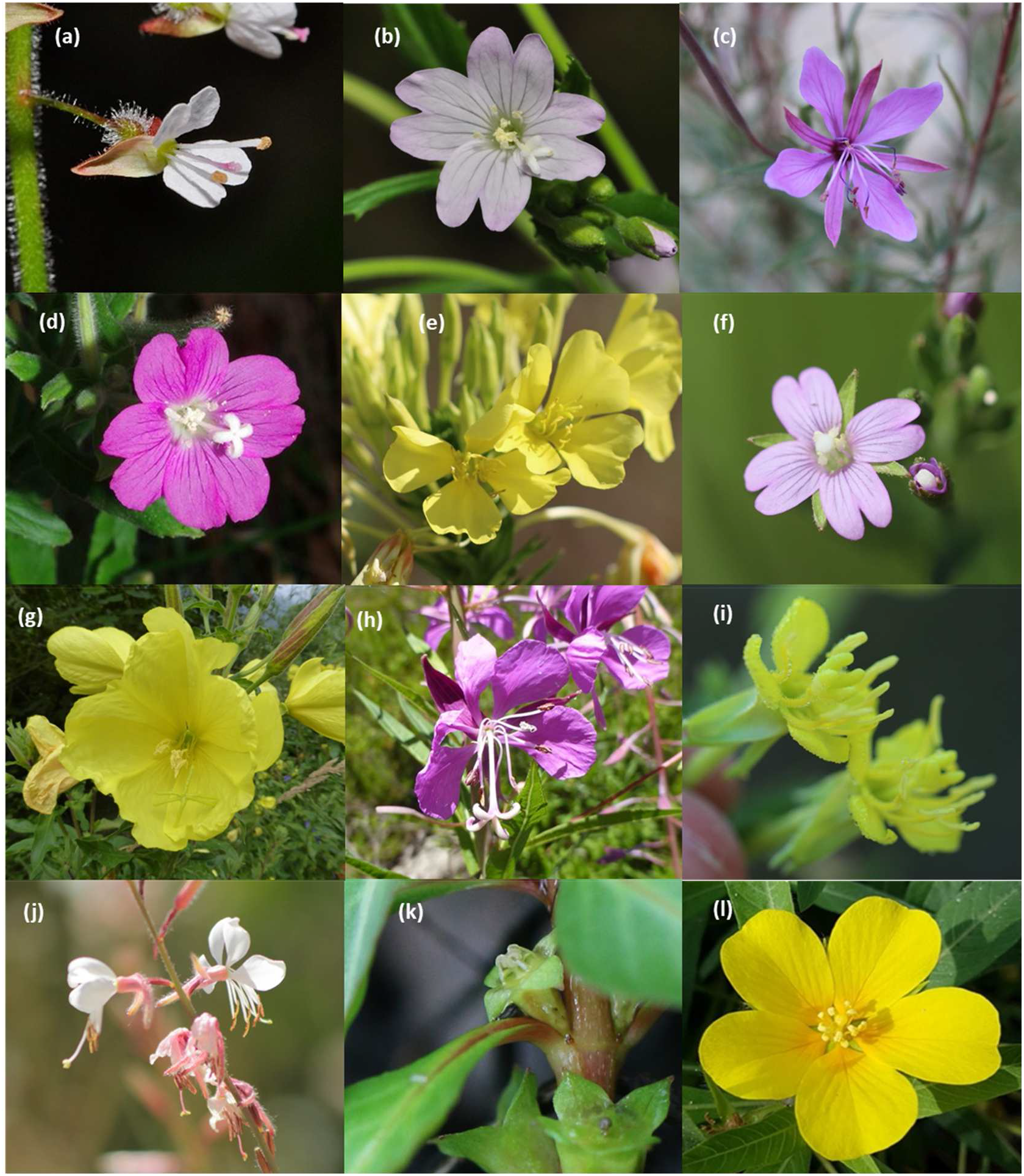
Floral diversity in Onagraceae family. (a) *Circaea lutetiana*. (b) *Epilobium montanum.* (c) *Epilobium dodonaei.* (d) *Epilobium hirsutum.* (e) *Oenothera pycnocarpa.* (f) *Epilobium tetragonum.* (g) *Oenothera glazioviana.* (h) *Chamerion angustifolium.* (i) *Oenothera subterminalis.* (j) *Gaura longiflora.* (k) *Ludwigia palustris.* (l) *Ludwigia grandiflora subsp hexapetala.* (a to k) images courtesy of Romain Deschamps, (l) image courtesy Luis Portillo.

**Figure S2:**
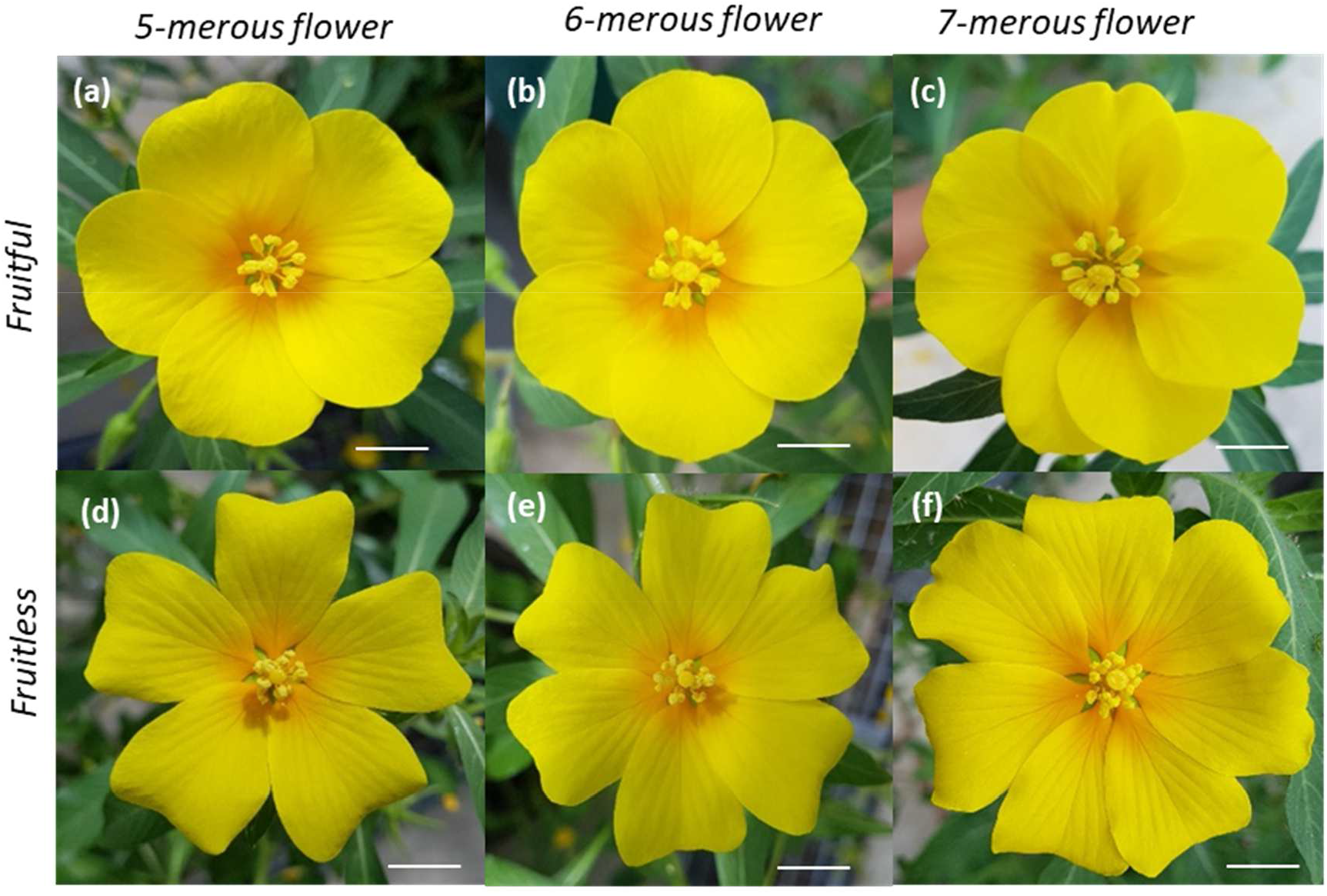
Floral morphology variation of *Ludwigia grandiflora* subsp *hexapetala* (2n=80): (a,b,c) 5-, 6- and 7-merous flowers from fruitful populations. (d,e,f) 5-, 6- and 7-merous flowers from fruitless populations. Floral formula for 5-merous flowers (a,d) were ⚥, Bt2, K5 + C5 + A10 + <u>G(5)</u>; for 6-merous flowers (b,e) were ⚥, Bt2, K6 + C6 + A12 + <u>G(6)</u> and 7-merous flowers (c,f) were ⚥, Bt2, K7 + C7 + A14 + <u>G(7)</u>. Bars = 1cm.

**Figure S3:**
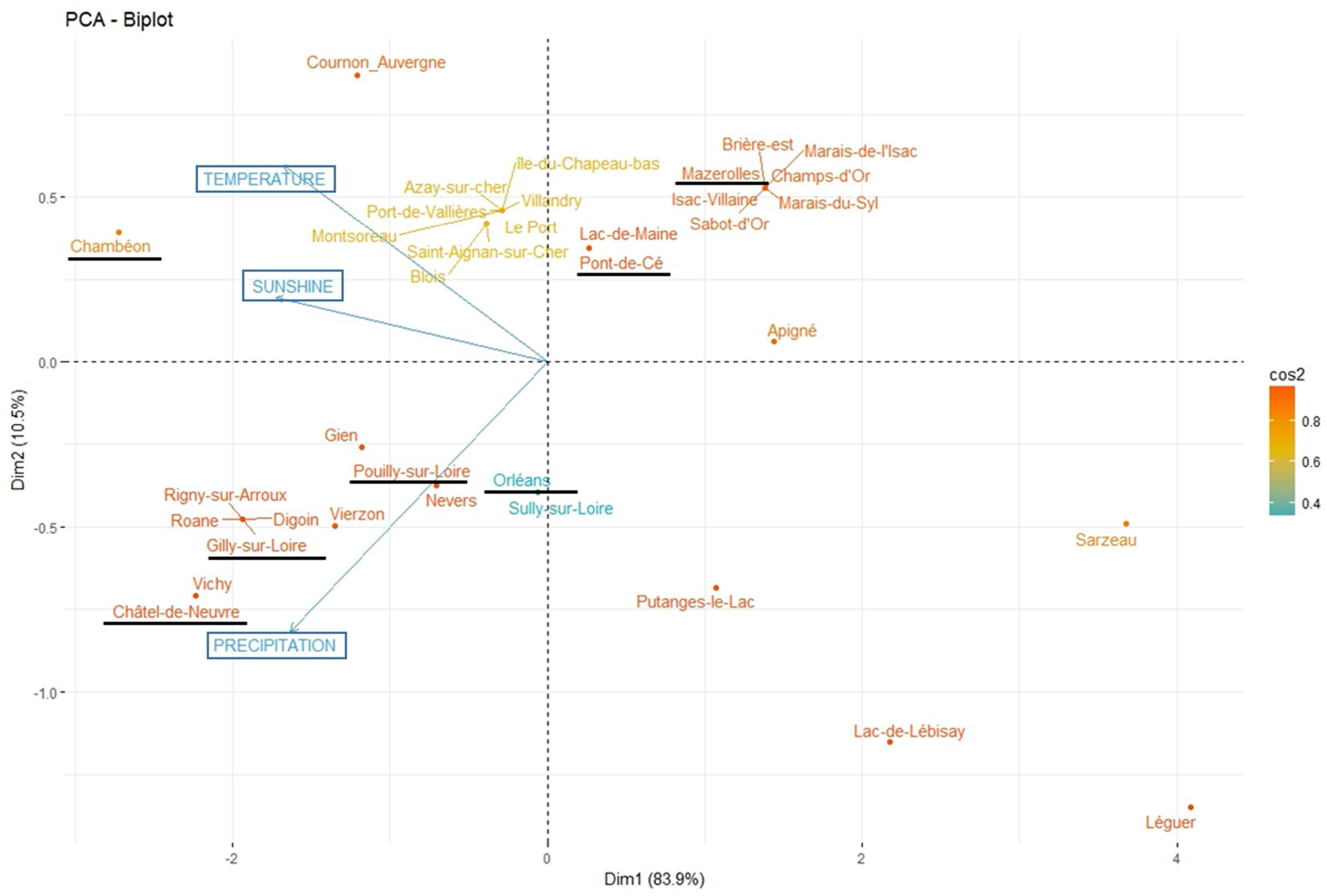
Principal Component Analysis of the distribution of *Ludwigia grandiflora* subsp. *hexapetala* populations in function of temperature, sunshine and precipitation in Loire basin. Colors (cos2) indicate variable contributions to axis, from low (blue) to high (red). The names of the 7 populations sampled for common garden and greenhouse experiments are underlined. Temperature was correlated to the sunshine and these two parameters inversely correlated to precipitation.

**Figure S4:**
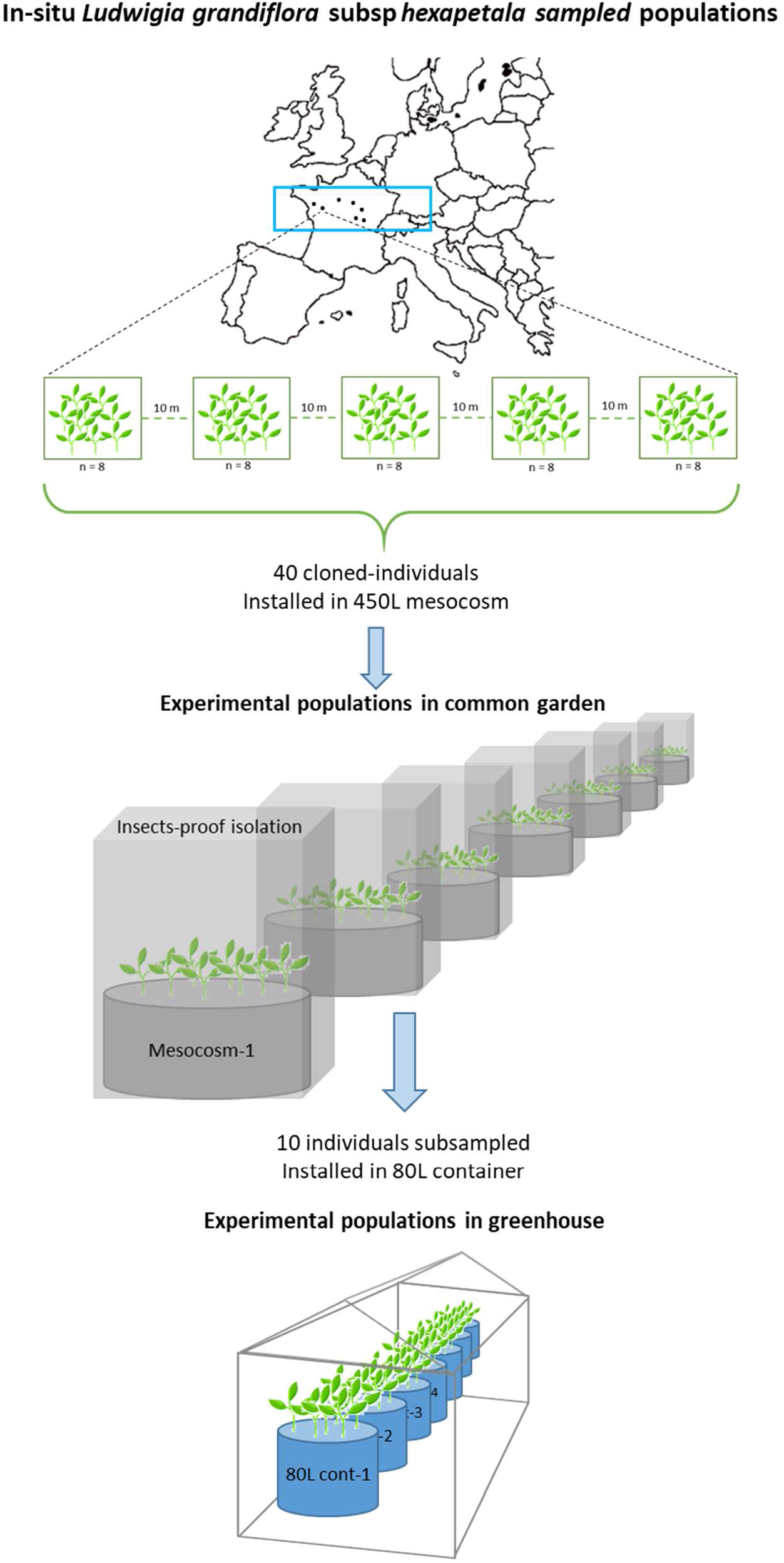
Sampling and experimental setup of *Ludwigia grandiflora* subsp. *hexapetala* populations *in-situ*, in common garden and greenhouse conditions. In the common garden, all 40 individuals sampled from populations bred together in a same mesocosm. In the greenhouse, all subsampled individuals were bred together in the same container.

**Figure S5:**
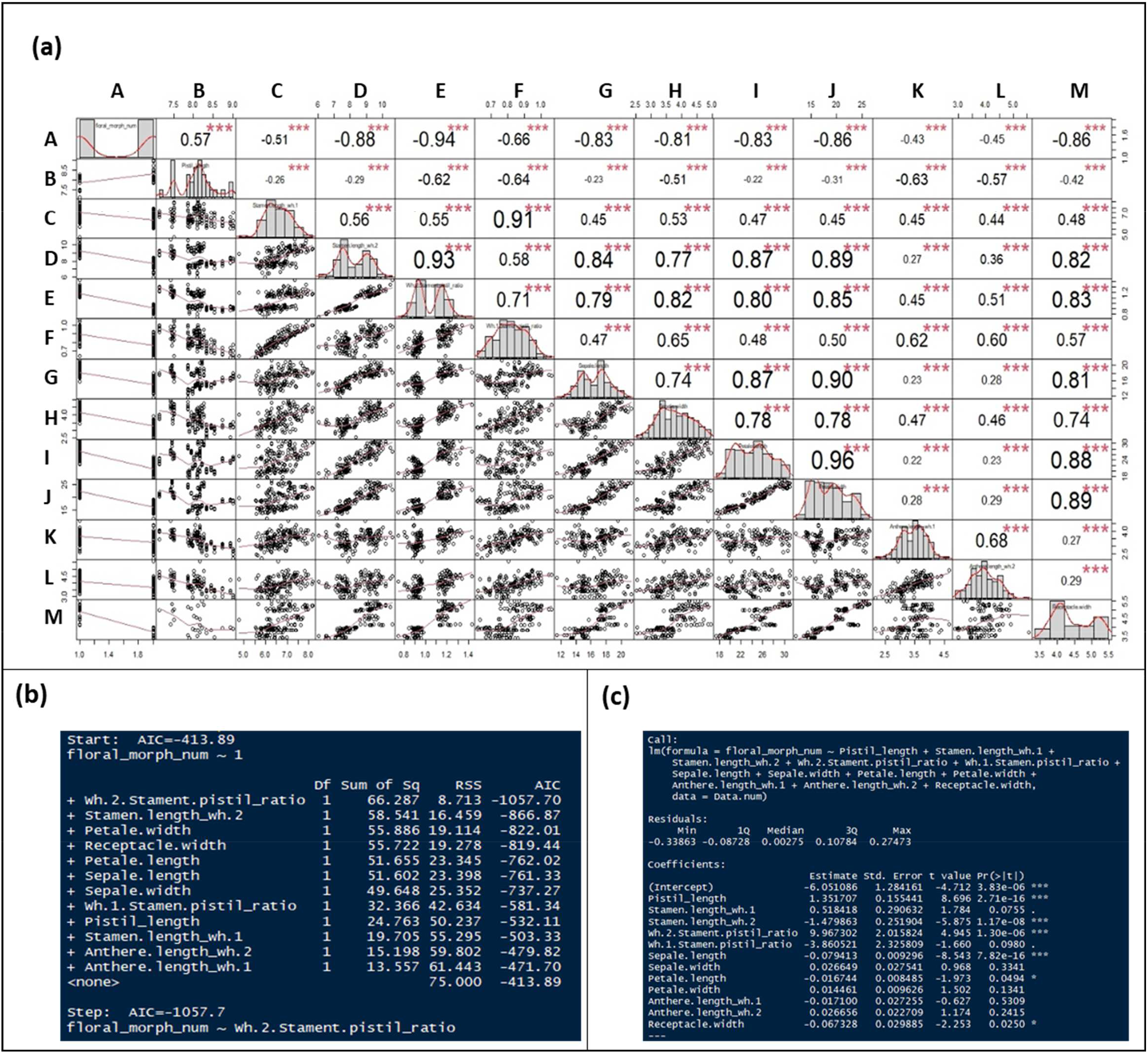
Multiple regression analysis of floral parts measurements. (a) Results of multiple correlation, A= floral morphs, B= pistil length, C= stamen length whorl-1, D= stamen length whorl-2, E= whorl-2 stamen-pistil ratio, F= whorl-1 stamen-pistil ratio, G= sepal length, H= sepal width, I= petal length, J= petal width, K= anther length whorl-1, L= anther length whorl-2, M= floral receptacle. (b) Model selection using Akaike Information Criterion (AIC) as a selection criterion. We obtained from the complete model’s (Floral_groups ~ 11 morphological traits) highest AIC value, AIC = −413.89, and the lowest AIC value from the most discriminant variable (floral_group ~ Wh.2.Stament.pistil_ratio) AIC= −1057.7 (C) Result of complete linear model analysis.

**Figure S6:**
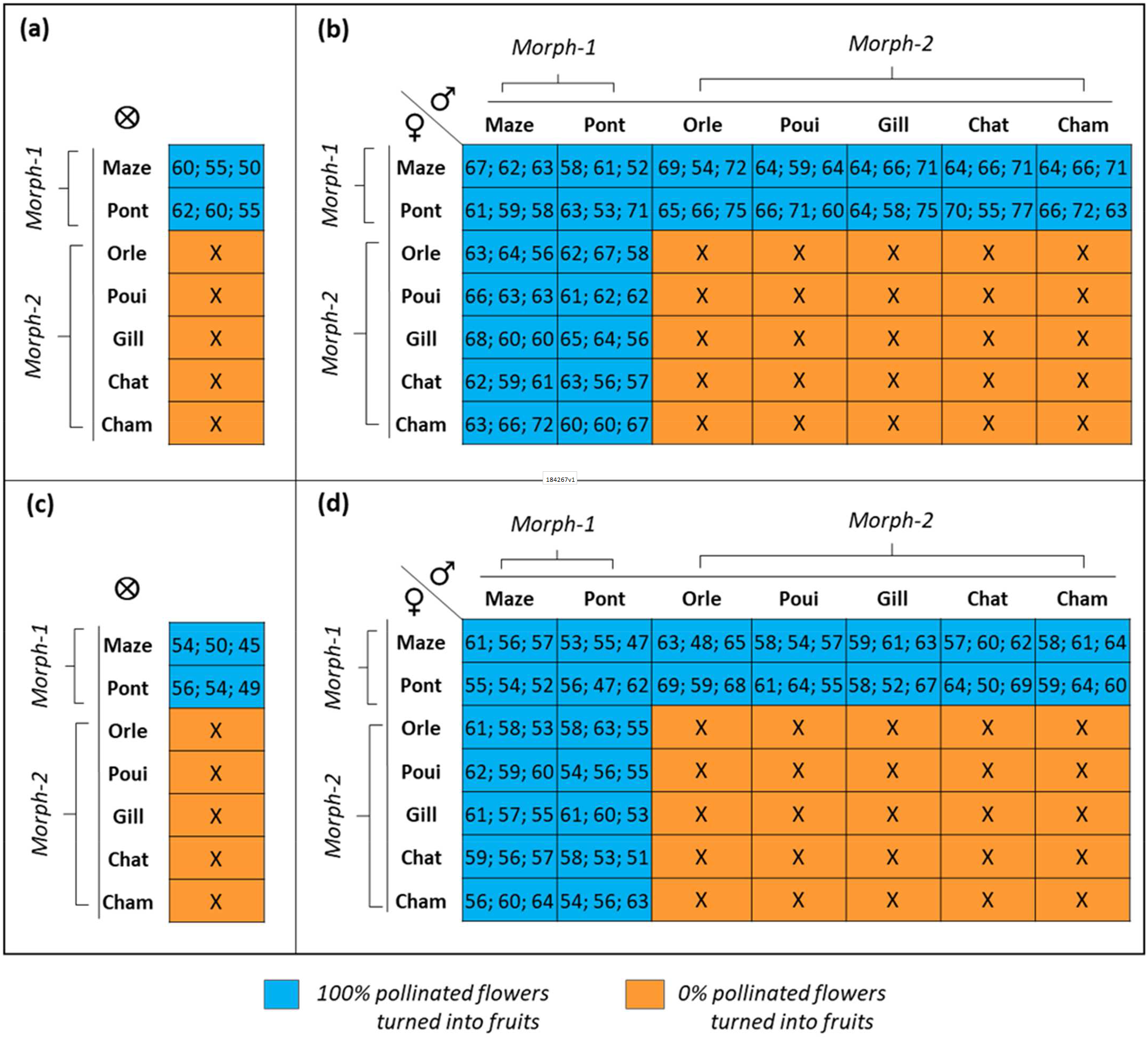
Fertility in the 7 sampled populations of *Ludwigia grandiflora* subsp. *hexapetala* after hand-controlled-pollination crosses from mid-July to beginning of August 2018 in greenhouse. (a) Seed production from 3 randomly selected fruits produced by self-pollination. (b) Seed production from 3 randomly selected fruits produced by cross-pollinations. In (a) and (b), numbers separated by semicolons stand for the number of seeds in fruit1, fruit2, fruit3. Cross sign indicates no fruit and no seeds. (c) Plant production from 3 randomly fruits produced by self-pollination. 61; 56; 57 = number of plants obtained from fruit1, fruit2, fruit3. Cross: no plant. (d) Plant production from 3 randomly selected fruits produced by cross-pollination. 61; 56; 57 = number of plants obtained from fruit1, fruit2, fruit3. Cross: no plant. In (c) and (d), numbers separated by semicolons stands for the number of fully-developed plant obtained from fruit1, fruit2, fruit3. Cross sign indicates no plant obtained.

**Figure S7:**
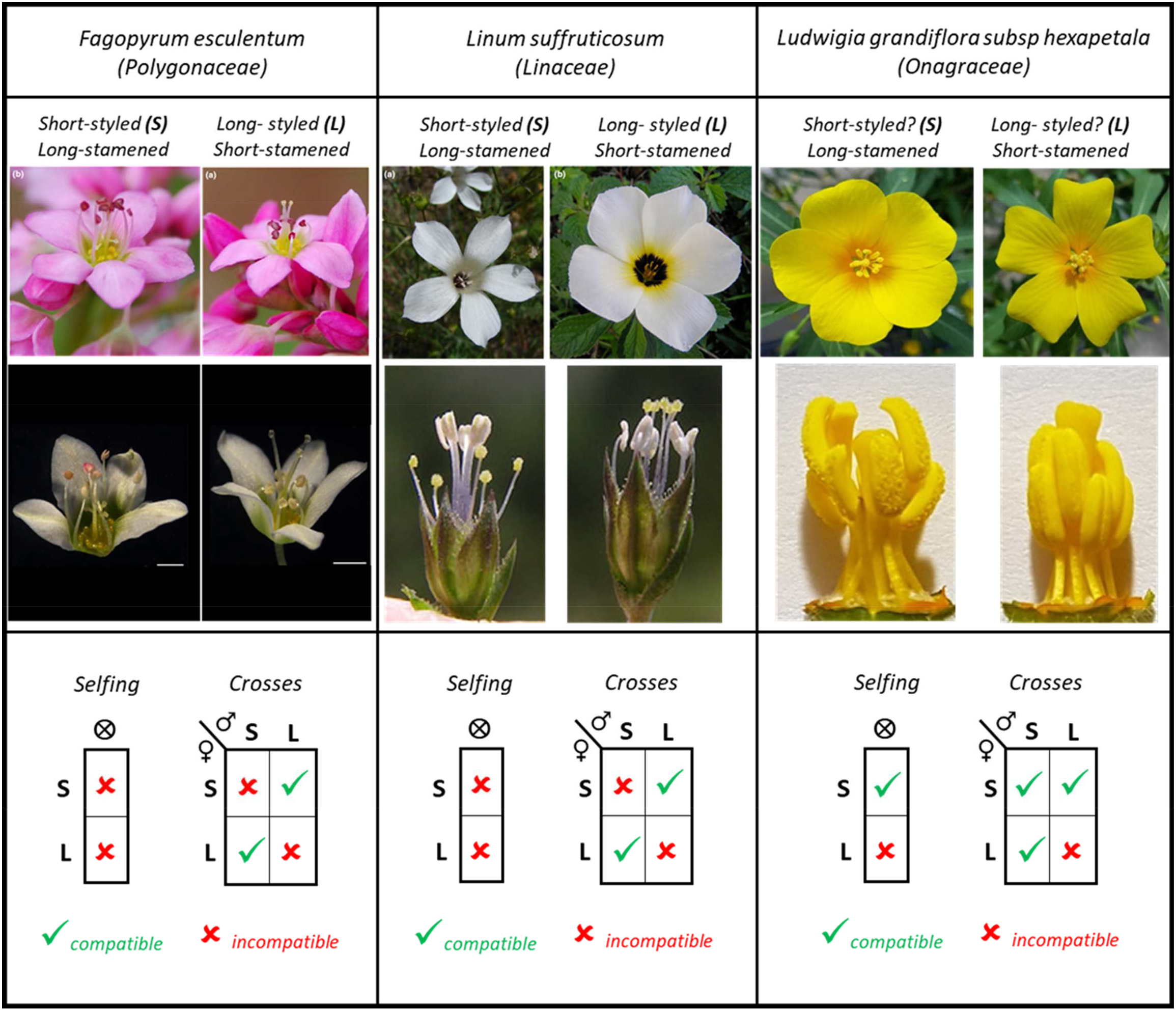
Floral morphology and reproductive system (compatible and incompatible cross) of 3 heteromorphic species: Left: *Fagopyrum esculentum* (pink flower from Barret *et al.* 2019; white flower from Li *et al*. 2017). Middle: *Linum suffruticosum* (*Barret et al.* 2019, Ruiz-Martín *et al.* 2018) Right: *Ludwigia grandiflora* subsp. *hexapetala* and mating schemas by Luis Portillo. “S” stands for short-styled flower, “L” for Long-styled flower. Green checked signs indicate fruitful and fertile crosses while red crosses indicate fruitless and infertile crosses.

**Table S1:**
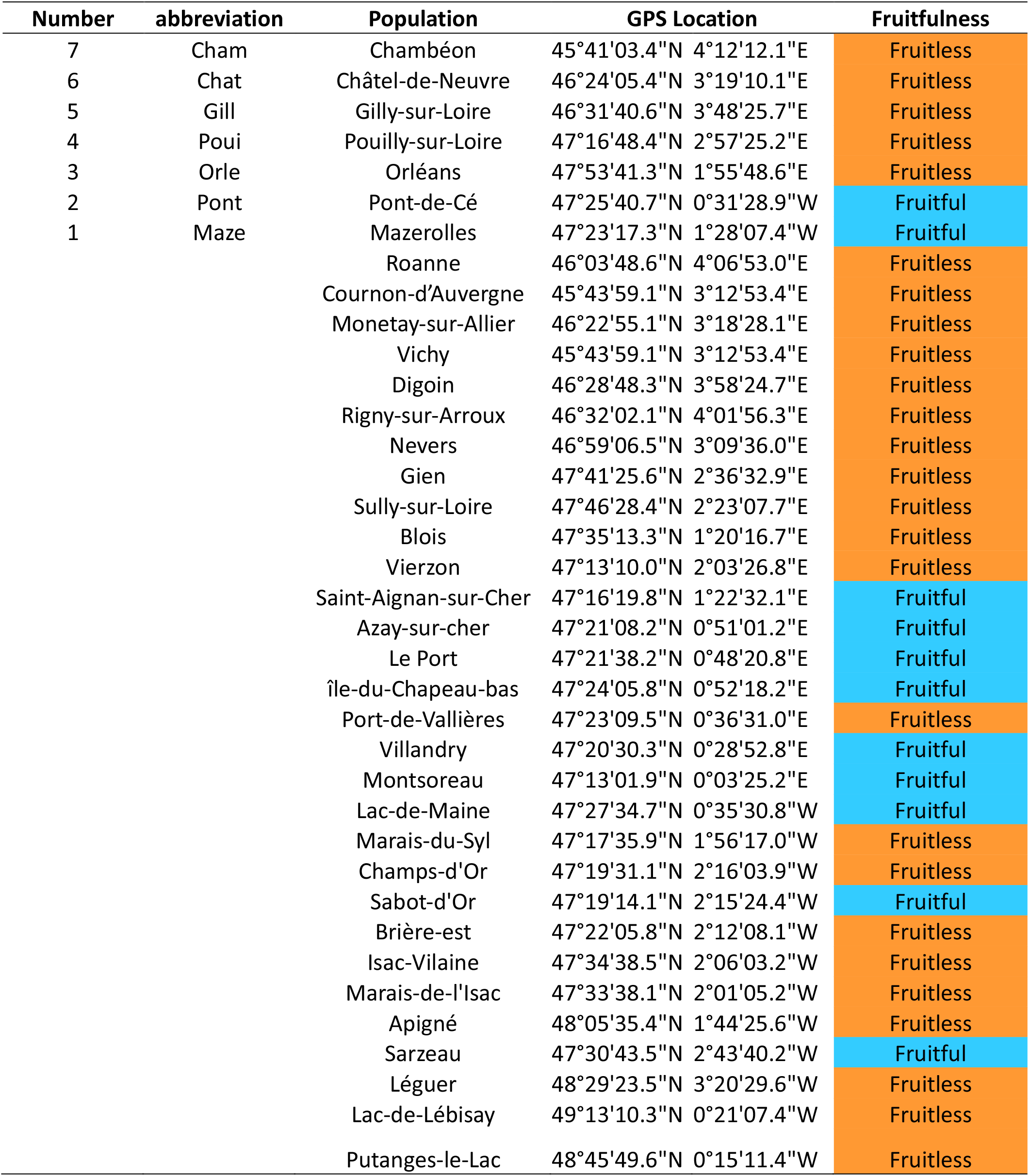
Location of all studied fruitful and fruitless populations in the Loire basin. The first seven populations were sampled for mesocosm and greenhouse experimentations.

**Table S2:** Mean values of floral parts biometrics data for floral group 1 and floral group 2 with their respective standard deviations.

| Floral parts | Floral group 1 | Floral group 2 |
| --- | --- | --- |
| Sepal length (mm) | 18.33 ± 1.12 | 14.91 ± 1.18 |
| Sepal width (mm) | 4.20 ± 0.36 | 3.28 ± 0.29 |
| Petal length (mm) | 27.15 ± 2.21 | 21.00 ± 1.91 |
| Petal width (mm) | 22.42 ± 2.06 | 16.26 ± 1.51 |
| First whorl stamen length (mm) | 6.97 ± 0.63 | 6.31 ± 0.47 |
| Second whorl stamen length (mm) | 9.31 ± 0.49 | 7.59 ± 0.43 |
| First whorl anther length (mm) | 4.12 ± 0.37 | 3.24 ± 0.44 |
| Second whorl anther length (mm) | 4.14 ± 0.41 | 3.68 ± 0.47 |
| Floral receptacle width (mm) | 4.87 ± 0.42 | 3.74 ± 0.2 |
| Pistil length (mm) | 7.82 ± 0.35 | 8.31 ± 0.35 |
| Nectar production (μL) | 9.43 ± 1.59 | 4.93 ± 1.99 |

**Table S3:**
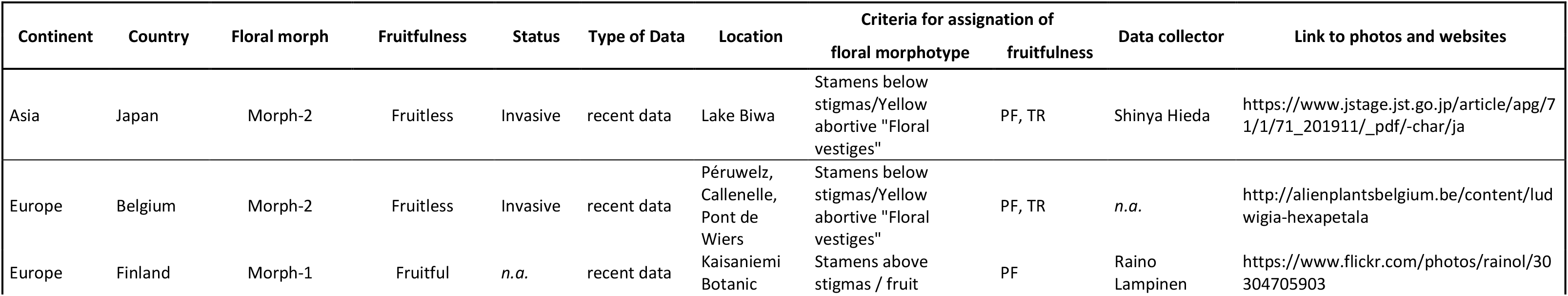
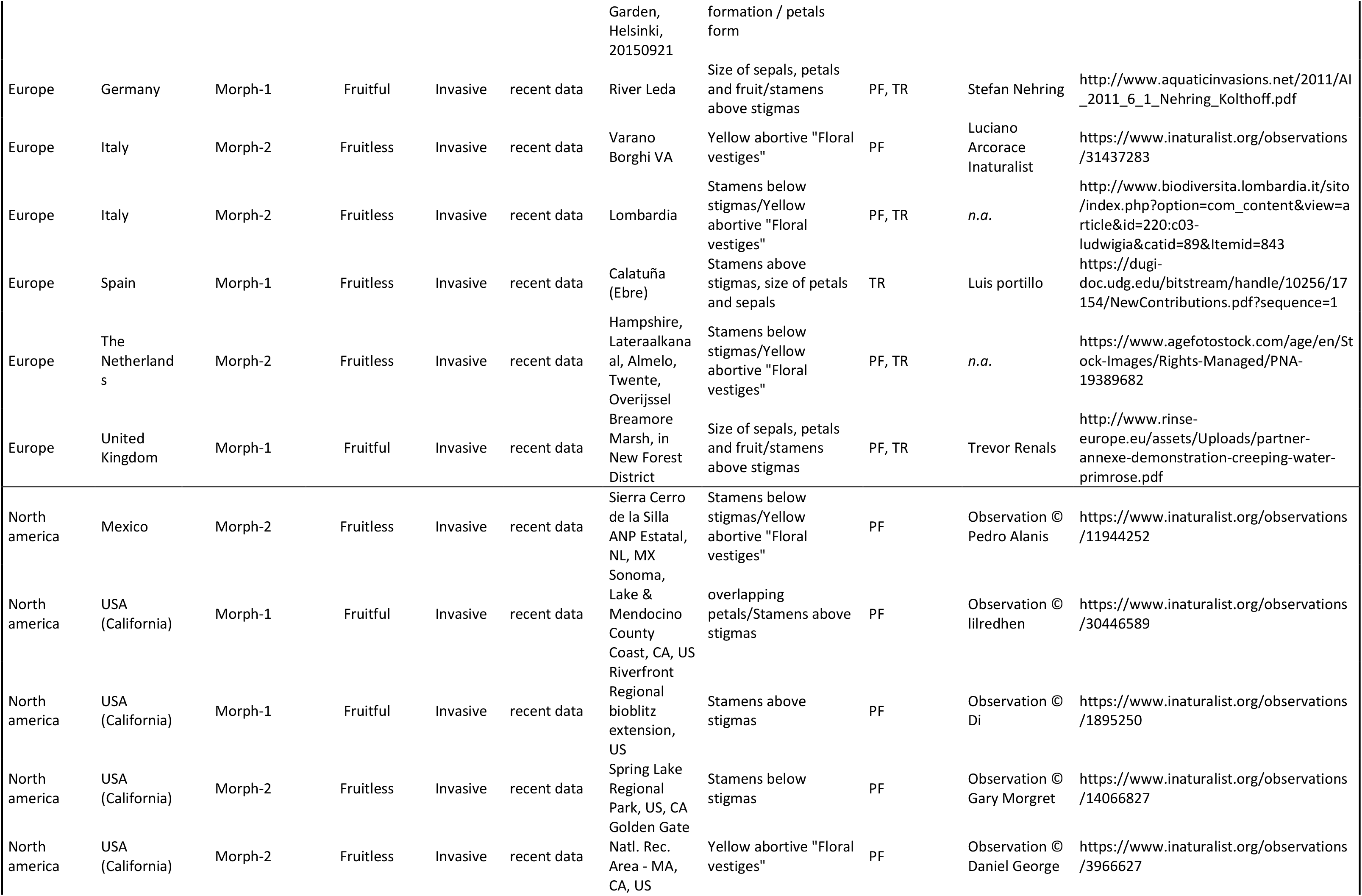
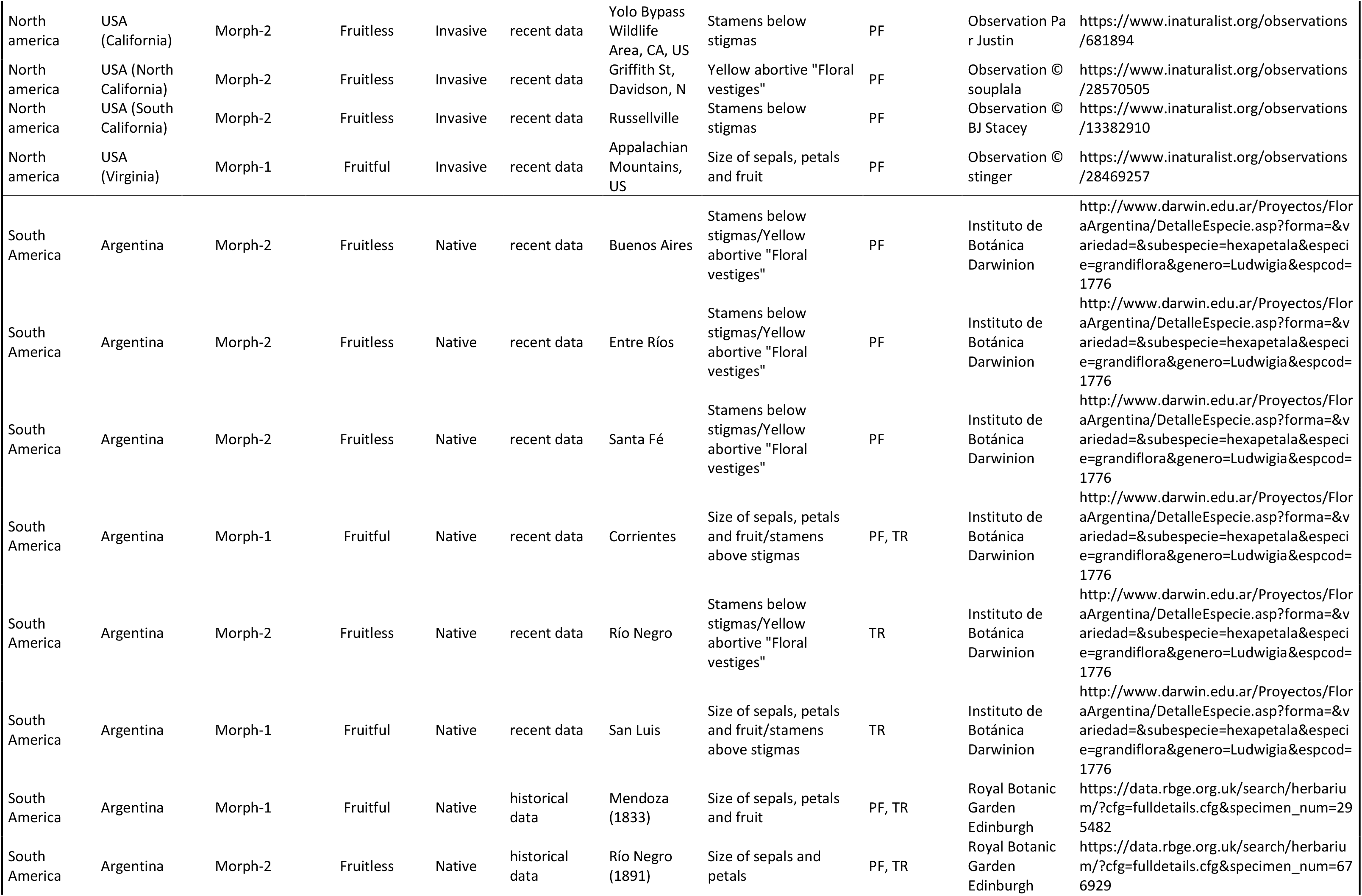
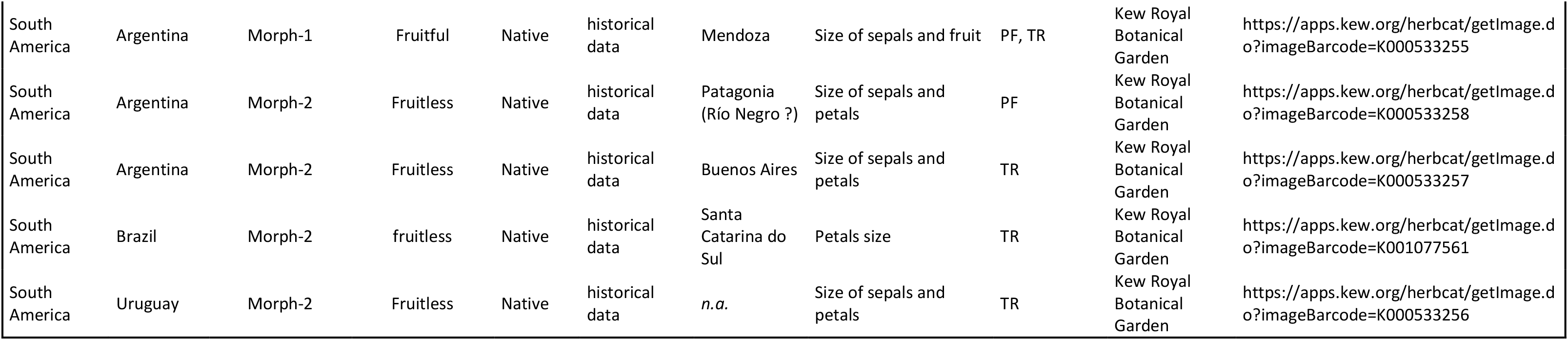
Assignment of floral morphs over the world using criteria defined for morphs observed in France, in *Ludwigia grandiflora* subsp. *hexapetala* (Syn. *Ludwigia hexapetala*) populations. Considering the binary answer we obtained in our results, we qualified fruitfulness in populations as ^**PF**^ when fruitfulness was directly observable from the available photos (PF) on which we saw either aborted fruits (fruitless) or fleshy developing fruit (fruitful) and ^**TR**^ using textual reports wrote by managers, naturalists and scientifics. Remark:

1. the list below included only images corresponding to the true species having a geographic origin, and showing at least one visible or measurable criterion of our biometrics.
2. Please note that many websites, even official and governmental ones, present significant and unfortunately frequent taxonomic errors. They annotated as *Ludwigia grandiflora* subsp. *Hexapetala* photos belonging to another *Ludwigia* species that can be distinguished by the shape of the leaves, presence / absence of trichomes on all plant organs, shape and size (length / width ratio) of the sepals, morphology of the lower ovaries, shape and size (length / width ratio) of fruits, deciduous or persistent sepals on fruit. Some examples:

i. https://bentonswcd.org/plant/large-flowered-primrose-willow/ Here photos correspond to *Ludwigia bonariensis*.
ii. https://www.inaturalist.org/observations/41006687 Here photos correspond to *Ludwigia leptocarpa*;
iii. https://www.inaturalist.org/observations/5994468; here photos correspond to *Ludwigia peploides* subs. *glabrescens.*
iv. https://www.inaturalist.org/observations/14123028 here photos correspond to hexaploid *Ludwigia grandiflora* subsp. *grandiflora* (syn. *Ludwigia grandiflora*).

## Notes

### Competing Interest Statement

The authors have declared no competing interest.

